# High-dimension to high-dimension screening for detecting genome-wide epigenetic regulators of gene expression

**DOI:** 10.1101/2022.02.21.481160

**Authors:** Hongjie Ke, Zhao Ren, Shuo Chen, George C Tseng, Jianfei Qi, Tianzhou Ma

## Abstract

**Motivation:** The advancement of high-throughput technology characterizes a wide range of epigenetic modifications across the genome involved in disease pathogenesis via regulating gene expression. The high-dimensionality of both epigenetic and gene expression data make it challenging to identify the important epigenetic regulators of genes. Conducting univariate test for each epigenetic-gene pair is subject to serious multiple comparison burden, and direct application of regularization methods to select epigenetic-gene pairs is computationally infeasible. Applying fast screening to reduce dimension first before regularization is more efficient and stable than applying regularization methods alone.

**Results:** We propose a novel screening method based on robust partial correlation to detect epigenetic regulators of gene expression over the whole genome, a problem that includes both high-dimensional predictors and high-dimensional responses. Compared to existing screening methods, our method is conceptually innovative that it reduces the dimension of both predictor and response, and screens at both node (epigenetic features or genes) and edge (epigenetic-gene pairs) levels. We develop data-driven procedures to determine the conditional sets and the optimal screening threshold, and implement a fast iterative algorithm. Simulations and two applications to long non-coding RNA and DNA methylation regulation in Kidney cancer and Glioblastoma Multiforme illustrate the validity and advantage of our method.

**Availability:** The R package, related source codes and real data sets used in this paper are provided at https://github.com/kehongjie/rPCor.

## 1 Introduction

Epigenetic alterations are inheritable traits that impact gene activity not involving changes in the DNA sequences (Gibney and Nolan, 2010). These modifications include a variety of molecular mechanisms such as DNA methylation, histone modification and noncoding RNAs and contribute to the pathogenesis of many diseases (Allis and Jenuwein, 2016). For example, epigenetic changes are present in all human cancers and considered one of the main factors in tumorigenesis and cancer development through altering expression of cancer related genes (Baylin and Jones, 2016). With the rapid development of high-throughput technology in the past decades, researchers now have access to a more comprehensive and accurate epigenome and capture a wide range of epigenetic modifications across the genome. Identifying epigenetic regulators of genes and investigating how they regulate the genes in disease associated pathways will provide mechanistic insights into the disease and have potential clinical usage. Compared to the *cis* (“local”) epigenetic regulators that are in physical promixity with the genes they act on, identifying the *trans* (“distal”) epigenetic regulators constitutes a more challenging problem since it involves assessment of a large number of potential regulatory relationships between epigenetic and gene markers over the whole genome. Novel statistical and computational approaches are needed to jointly analyze these high-dimensional omics data and reveal the complex relationship between epigenetics and gene expression.

A widely used procedure that targets at such problems involving a large number of responses (genes) and a large number of predictors (epigenetic features) is to test the association for each epigenetic-gene pair separately. A canonical example is the association analysis between DNA variants and gene expression known as expression quantitative trait locus (eQTL) analysis (Aguet and Muñoz Aguirre, 2017). However, such a univariate analysis approach faces very serious multiple comparison issue and tends to be over-conservative after multiple testing correction. An alternative procedure is to fit a multivariate regression model to accommodate all predictors and responses and then perform model selection using regularization methods. In epigenetic-gene regulation problem, however, both predictor and response spaces are of extremely high dimension, directly rendering any regularization methods in this model becomes computationally infeasible. One common remedy for such computational bottleneck is to apply dimension reduction techniques such as sure screening methods first to quickly reduce the dimension without incorrectly removing the true signal pairs, then apply regularization methods for the final model selection in a lower dimensional space (Fig 1(a), red edges regarded as true signal pairs). Such a two-stage procedure (screening + regularization) is shown to be more efficient and stable than regularization methods alone and has wide applicability for big data analysis with desired scalability and theoretical guarantees. Since Fan and Lv (2008) proposed the first sure screening method “SIS” using the marginal correlation, numerous screening methods have been developed for dimension reduction in different types of models and using different statistics (Fan and Song, 2010; Li et al., 2012; Liu et al., 2015). Nevertheless, these screening methods usually look at univariate response, thus they are not directly applicable to the epigenetic regulation problem.

**Figure 1:**
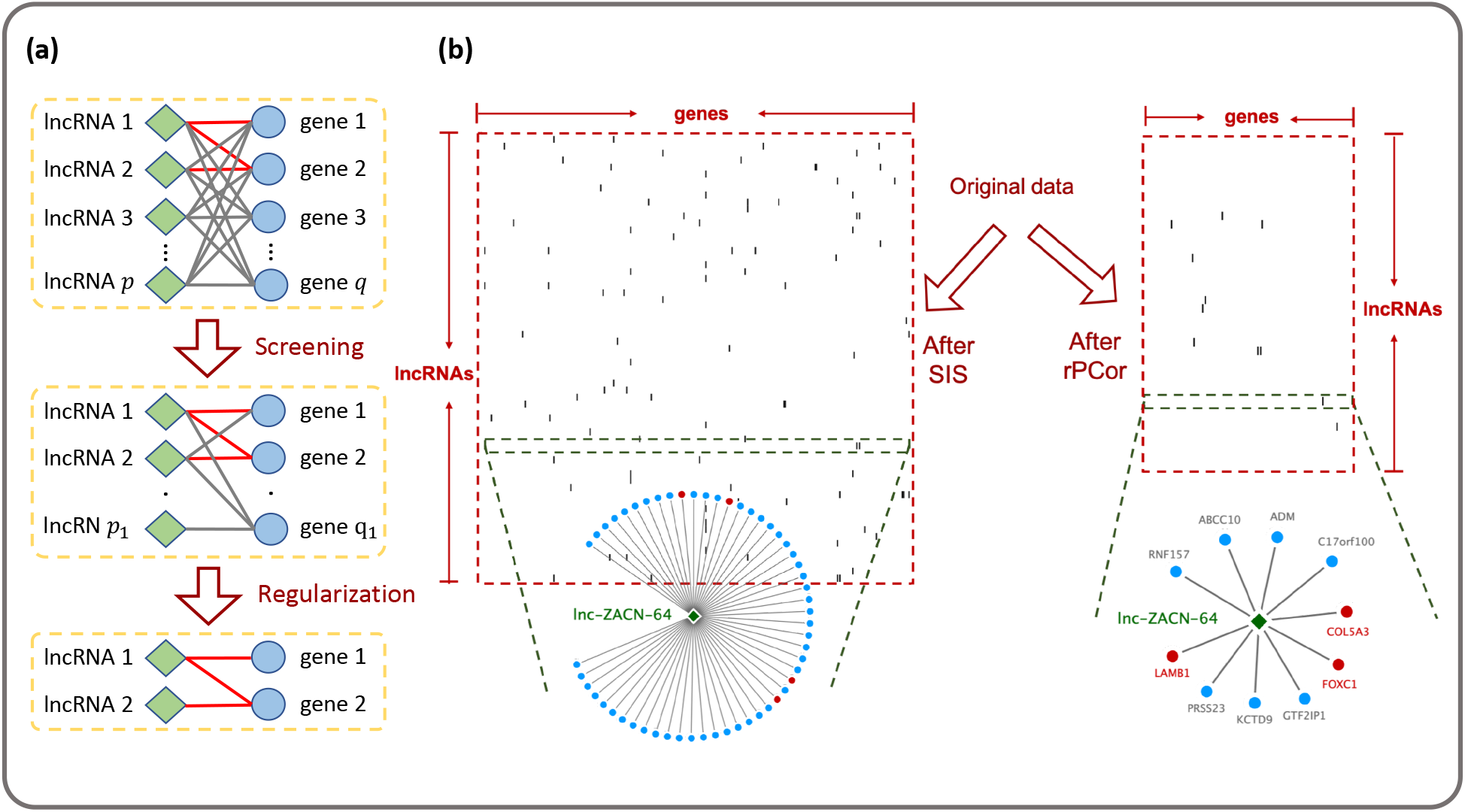
(a) Concept of the two-stage procedure (screening + regularization) in epigenetic-gene regulation using lncRNA regulation as an example. Screening stage will reduce the dimension to a more manageable scale (*p*_1_ ≪ *p*, *q*_1_ ≪ *q* and much fewer edges) without removing the true signals (red edges refer to the underlying true epigenetic-gene regulatory pairs). (b) lncRNA-gene regulation example in real data application to demonstrate the power of our screening method as compared to the SIS. The original data has 2,170 lncRNAs and 13,408 genes. Only 3% of lncRNAs and genes are shown here. After screening by SIS (left), 2,170 lncRNAs, 13,406 genes and 9,715,838 pairs are left, the matrix of remaining pairs (black dotted) still looks very dense. Each lncRNA (using “lnc-ZACN-64” as an example, the row highlighted in green) still has many remaining genes which can be potentially false positives. After screening by our method “rPCor” (right), only 1,652 lncRNAs, 5,274 genes and 7,528 pairs are left, the matrix of remaining pairs looks very sparse. Each lncRNA has much fewer number of genes remained, but the most important genes in the top disease related pathways (red colored, regarded as true signals) are kept.

A naive way to handle multiple responses is to implement screening for each response separately. This can be unsatisfactory by ignoring the complex correlations between responses and still leaves us with a large model. Figure 1(b) shows part of data from the long noncoding RNA (lncRNA) regulation example in Section 4. When SIS was applied separately for each gene (left), the matrix of remaining lncRNA-gene pairs (black dotted) looks very dense. Almost no lncRNAs or genes are removed thus no dimension reduction is achieved. Each lncRNA (using “lnc-ZACN-64” as a representative) has hundreds of gene targets left after SIS, many of which can be potential false positives. When our method “rPCor” was applied (right), the matrix of remaining lncRNA-gene pairs becomes much more sparse. Our method manages to remove lncRNAs with no target genes and genes not regulated by any lncRNAs, reducing the dimension of both predictor and response spaces. There are much fewer genes left for each lncRNA but almost all important gene targets in disease related pathways (red colored, regarded as true signals) remain.

In this paper, we propose a novel high-dimension to high-dimension screening method based on robust partial correlation, namely “rPCor”, for detecting epigenetic regulators of gene expression over the whole genome. To the best of our knowledge, our method is the first screening method that targets at a problem with both high-dimensional predictors and high-dimensional responses. Comparing to existing screening methods, our method is conceptually innovative that it can reduce the dimension of both predictor and response, and screens out both irrelevant nodes (epigenetic features or genes) and edges (epigenetic-gene pairs). We develop data-driven procedures to determine the conditional set and the optimal screening threshold and implement an iterative algorithm which is computationally feasible with hundreds of thousands of predictors and responses. The tail robustified partial correlation is used to protect against non-normality and heavy-tailed distributions to achieve the best screening performance. Extensive simulations and two real data applications in lncRNA and DNA methylation regulation of gene expression in cancers using The Cancer Genome Atlas (TCGA) data have shown the significant advantages of our method and its potential to be widely applied as a first stage pass for similar big data integration problems. The rest of the paper is organized as follows. In Section 2, we first introduce the model and the main characteristics of our screening method. In Section 3 and 4, we present the simulation studies and real data application examples. Final discussion is provided in Section 5.

## 2 Methods

### 2.1 Model and review of existing methods

Suppose we have collected both gene expression (e.g. from microarray study or RNA-seq study) and epigenetic data (e.g. DNA methylation from array or bisulfite sequencing) over the whole genome for *n* subjects. To model the dependency of gene expression on epigenetics and identify the important epigenetic regulators of genes, we consider a multivariate regression model with genes being the responses and epigenetic features being the predictors:

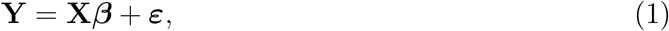

where 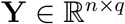 is the response matrix of expression of *q* genes for *n* subjects, 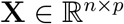 is the predictor matrix of values of *p* epigenetic features for *n* subjects, and 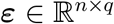 refers to the random error terms that have a joint multivariate distribution with mean **0** and covariance matrix **Σ_*Y*_**. Here we consider **X** consists of one type of epigenetic data for simplicity, though the same regression framework also applies to modeling the additive effects of mixed types of epigenetic data.

Our purpose is to identify the nonzero coefficients in matrix 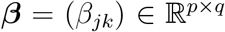, i.e. the set

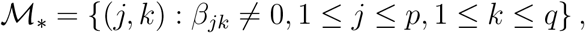

which consists of important epigenetic-gene regulatory pairs. In low dimensional case where *p* and *q* are smaller than *n*, the ordinary least square (OLS) estimation can be used to estimate the coefficients. When the numbers of predictors and responses grow to much larger than the sample size (i.e. *p* >> *n, q* >> *n*) as in epigenetic-gene regulation problem, classical OLS estimation fails and we usually need the sparsity assumption to tackle the problem. For univariate response (when *q* = 1), it is common to assume only a small number of predictors are relevant with nonzero coefficients, selecting predictors is equivalent to selecting predictor-response pairs. For multiple response, however, selecting nodes (predictors or responses) is not equivalent to selecting edges (predictor-response pairs). Thus, we further consider sparsity at two levels: (1) overall sparsity or edge-wise sparsity in ***β***, i.e. a majority of *β_jk_* = 0; (2) column-wise/row-wise sparsity or node-wise sparsity, i.e. *β_jk_* = 0 for the entire row or column. In real data when we investigate the epigenetic regulation for a particular disease, it is natural to assume that only a few true epigenetic-gene regulatory pairs exist and these pairs are restricted to a small set of critical epigenetic biomarkers (e.g. master predictors) which regulate a small set of responsive genes (e.g. key responses), while others are not participating in the relevant regulatory process (Cheng et al., 2019).

Direct regularization methods that enforce sparsity at both edge and node levels have been developed for the multivariate regression model. Peng et al. (2010) adopted a mixed *ℓ*_1_/*ℓ*_2_ norm penalty in searching for master predictors that affect many responses. Li et al. (2015b) proposed a similar but more general multivariate sparse group lasso to encourage the selection of both individual and grouped coefficients in ***β*** (e.g. treating coefficients of all responses to the same predictor as a group). The *ℓ*_2_ penalty shrinks unimportant predictors with small aggregate effects over all responses to zero, however, when *q* is large, almost all predictors will have large aggregate effects thus no master predictors are prioritized. In addition, when the product *pq* is extremely large, these regularization methods usually become computationally intractable and unstable.

An alternative is a two-stage strategy where we first apply sure screening methods to reduce the dimension of the model, and then apply regularization methods to a model with lower dimension for the final selection. Fan and Lv (2008) proposed the first sure screening method called Sure Independence Screening (SIS) that only kept the predictors with large marginal correlation with the response in univariate linear model and showed its sure screening property that the set after screening will contain the true set of predictors with large probability. Alternative marginal association based screening methods, including those adopting robust and model-free statistics, were developed and generalized to a broader class of models (Fan and Song, 2010; Zhu et al., 2011; Ma et al., 2020). Other screening methods have been proposed to further consider the between-predictor correlation (Bühlmann et al., 2010; He et al., 2019). See Supplement section 1 for a selective overview of the popular screening methods. However, all these screening methods mainly consider univariate response thus do not directly apply to multivariate regression model.

For the model with multiple responses, one can apply existing screening methods for univariate response to each response to select the top predictor-response pairs, however, this approach ignores the correlation between responses and potentially generates many false positives. In addition, applying screening to each response tends to ignore the nodewise sparsity and leaves almost all responses in the model after screening. Some screening procedures that can handle multiple responses (Li et al., 2012; Di He and Zou, 2021) used statistics that aggregates over all responses. These methods work well in response spaces with low dimension. However, when *q* >> *n*, they face the same challenges as the aforementioned regularization methods that almost all predictors have large aggregate effects, thus can hardly achieve dimension reduction.

We propose a robust Partial Correlation based method, namely rPCor, to perform screening for models with high-dimensional predictors and high-dimensional responses. The purpose of our method is to screen out as many noisy predictor-response pairs while keeping most of the predictor-response pairs from the true model. After screening, we apply regularization method to refine the final pool and select the most important pairs. Figure 2 shows a flowchart of the two-stage strategy for epigenetic-gene regulatory pair selection. The rPCor based screening stage consists of three steps that will reduce the original dimension *p* × *q* to a more manageable scale *p*_1_ × *q*_1_ (i.e. only *p*_1_ predictors and *q*_1_ responses are left where *p*_1_ < *p*, *q*_1_ < *q* and other nodes have zero *β*’s for the entire row or column). Screening of edges is indicated by the sparsity indicator matrix **U**_*P*_1_×*q*_1__, where = *U_lh_* if the (*l, h*) pair is kept and *U_lh_* = 0 if it is removed after screening. The sparsity indicator matrix **U** will later serve as input for the regularization stage to get the final *β* estimate.

**Figure 2:**
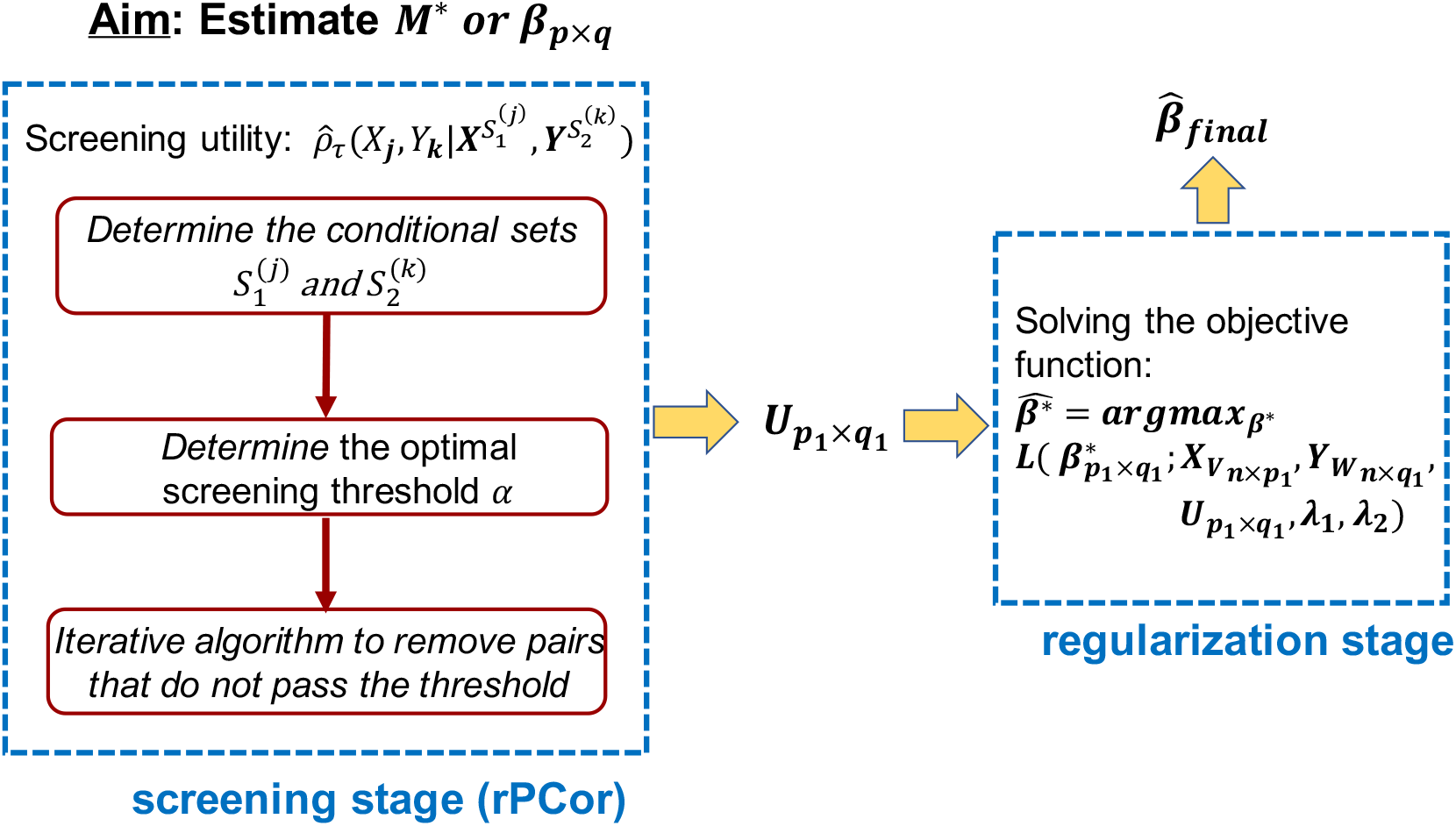
Flowchart of the two-stage (screening+regularization) strategy for epigenetic-gene regulatory pair selection (i.e. estimating 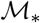 or ***β_p×q_***). Screening stage by rPCor will reduce the dimension of *p* × *q* to a more manageable scale *p*_1_ × *q*_1_ (i.e. only *p*_1_ predictors and *q*_1_ responses are left where *p*_1_ < *p*, *q*_1_ < *q*). Screening of edges is indicated by the sparsity indicator matrix **U**_*p*_1_×*q*_1__, where *U_lh_* = 1 if the (*l,h*) pair is kept and *U_lh_* = 0 if it is removed after screening, and will be used as input in the regularization stage for final *β* estimation.

### 2.2 Robust partial correlation based screening

The relationship between partial correlation and regression coefficients has been widely studied before (Peng et al., 2009). Bühlmann et al. (2010) proposed to use the partial correlation 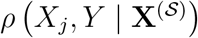, *j* = 1,…, *p* for screening in linear model and made the following partial faithfulness assumption to theoretically justify its use (see Supplement section 2 for more details on this assumption and the connection between partial correlation and regression coefficient): 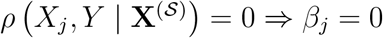, where *S* is some subset of {*j*}^*C*^.

For univariate response, only the conditional set for predictors needs to be included. For multiple responses, in order to adjust the inter-feature correlations among responses, we include both predictors and responses in the conditional sets, which helps remove more false positive edges and nodes. More specifically, we propose to use the partial correlation 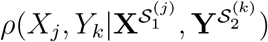, *j* = 1,…, *p*; *k* = 1,…, *q* for screening in multivariate regression model, where 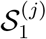 and 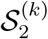 are the conditional sets for the nodes *X_j_* and *Y_k_*, respectively. We develop procedures to carefully determine the conditional sets for each node which greatly improve the computation and screening performance. We impose the following extended partial faithfulness assumption to justify its use:

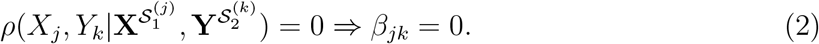

Based on the extended partial faithfulness assumption, we want to keep the following set:

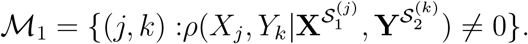

In finite samples, one can estimate the sample partial correlation to obtain the above set as long as the cardinality of the conditional set is not too large. However, the sample partial correlation estimate is not robust against heavy-tailness, a notable feature of highdimensional data that has brought new challenges to many statistical methods. Although proper transformation of expression data (log-transformation) and methylation data (M-values) are commonly used to better fit the normality assumption, heavy-tailness and outliers are still inevitable in real data. To protect against the violation of normality assumption and heavy-tailness in both epigenetic and gene expression data, we propose to estimate a tail robustified version of the partial correlation:

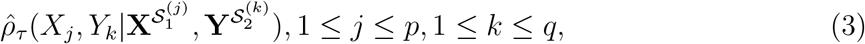

where 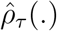 is the partial correlation estimate after applying the truncation operator *ψ_τ_*(*x*) = (|*x*| Λ *τ*)sign(*x*) to both *X_j_* and *Y_k_* adjusting for the variables in the conditional sets 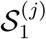 and 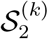, |*x*| Λ *τ* = min{|*x*|, *τ*}, and *τ* > 0 is called the robustification parameter. We follow from the literature (Ke et al., 2019) and propose to adopt a data-adaptive approach to automatically tune *τ* and obtain the robustified estimate for each partial correlation computed (see Supplement section 3). We then apply Fisher’s *Z* transform to the robust partial correlation estimate to obtain 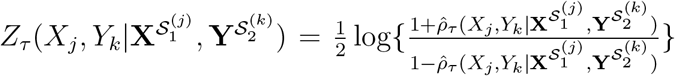 and identify the following set as an estimate of the set 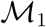:

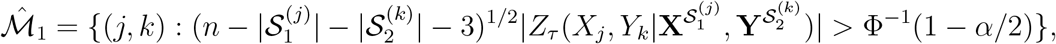

where Φ is the cumulative density function of standard normal distribution and |.| is the cardinality of the set. Here *α* is the screening threshold parameter that determines how aggressive or conservative users want to be at the screening stage. Some methods (Fan and Lv, 2008; Li et al., 2012) directly tune the model size after screening with theoretical guidance, which served the same role as *α* in practice. The choice of *α* is critical to screening methods that guarantees the set after screening includes as fewer false positives as possible while retaining all true positives. We propose a stability selection based procedure to determine the optimal *α* in practice as described next.

### 2.3 Conditional set and threshold parameter selection

Partial correlation based methods typically search over all nodes in the full conditional set when testing for the partial correlation, which is computationally heavy. In epigenetic-gene regulation problem, it is natural to assume that a majority of epigenetic features (predictors) are unrelated and most genes (responses) are unrelated over the whole genome. When we compute the partial correlation, we can restrict the conditional sets to only neighboring nodes for each node of interest, which will greatly improve the computation and the screening performance. To determine the conditional sets for both epigenetic predictors (i.e. 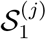) and gene responses (i.e. 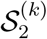), we propose to apply a marginal correlation threshold and regard only those predictors/responses having marginal correlation above certain threshold with the predictor/response of interest as the conditional sets, i.e. 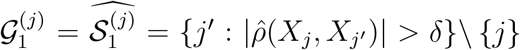 and 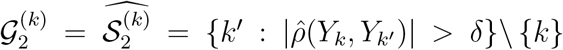 where 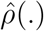 refers to the sample marginal correlation. One may also use a tail-robustified version of marginal correlation in the above calculation. For the cutoff *δ*, we randomly generate variables from two independent distributions to calculate spurious correlations under the same sample size and set *δ* to be the maximum spurious correlation among multiple replications. Given the fact that module structure naturally exists in gene expression data, we also provide an alternative solution to determine the conditional sets by first partitioning genes into unconnected modules and then only include genes within the same module in the conditional set (see Supplement section 4).

The choice of screening threshold parameter *α* is critical to all screening methods as it balances between the sensitivity and false positive control of the method. If *α* is chosen too small (too stringent cutoff), it is possible that important signal pairs will be accidentally screened out in the screening stage. On the other hand, if *α* is chosen too large (too loose cutoff), most pairs are kept so the model remains large which makes the screening step meaningless. Most screening methods provide theoretically guided choice of *α* without giving empirical guidance on how to find its optimal value in practice. To determine a data-driven choice of *α*, we propose a stability selection based procedure (Meinshausen and Bühlmann, 2010). The idea of stability selection is to identify variables that are included in the model with high probabilities when a variable selection procedure is performed on random samples of the observations. In our case, the optimal *α* is chosen such that the selection frequencies of top predictor-response pairs among random subsamples are large enough to pass a threshold determined by controlling the empirical Bayes false discovery rate. We provide details of this procedure in Supplement section 5 as well as a figure to illustrate how we apply this procedure to select *α* = 1*e* — 5 for TCGA GBM real data example (Figure S1).

Table 1 lists the main characteristics of our method “rPCor” and its comparison with other popular state-of-the-art screening methods. Compared to the marginal or partial correlation based screening methods in univariate model, we further consider conditioning on both predictors and responses to reduce false positives. We also propose data-driven procedures to determine the conditional sets and the optimal screening threshold to improve screening performance and computation efficiency. Lastly, our method is robust against heavy-tailed distributions.

**Table 1:**
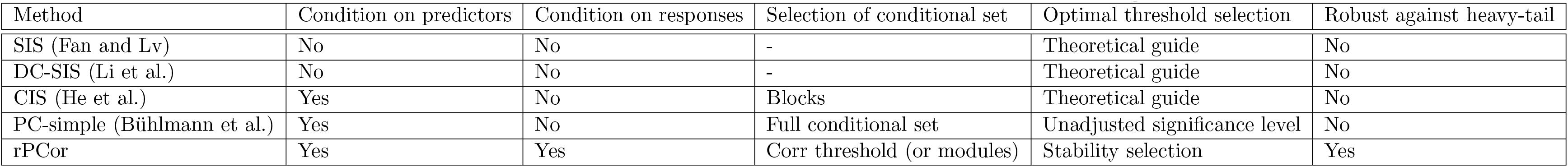
A comparison of our method with other popular state-of-the-art screening methods

| Method | Condition on predictors | Condition on responses | Selection of conditional set | Optimal threshold selection | Robust against heavy-tail |
| --- | --- | --- | --- | --- | --- |
| SIS (Fan and Lv) | No | No | - | Theoretical guide | No |
| DC-SIS (Li et al.) | No | No | - | Theoretical guide | No |
| CIS (He et al.) | Yes | No | Blocks | Theoretical guide | No |
| PC-simple (Bühlmann et al.) | Yes | No | Full conditional set | Unadjusted significance level | No |
| rPCor | Yes | Yes | Corr threshold (or modules) | Stability selection | Yes |

### 2.4 Iterative algorithms

Directly computing the partial correlation can still be burdensome in spite of pre-determination of the conditional sets 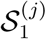 and 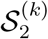. We follow the same idea of PC-simple algorithm (Bühlmann et al., 2010) and propose a fast iterative algorithm by starting with marginal screening (zero order) and sequentially increasing the order of partial correlation until the size of the active set is no greater than the order or certain maximum order is reached (see Algorithm 1).

### 2.5 Regularization after screening

After screening, the dimension is reduced to a more manageable scale. Define the node subsets 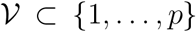 with 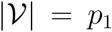 and 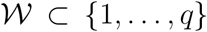 with 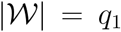, i.e. only *p*_1_ epigenetic features and *q*_1_ genes are left after screening. The predictor matrix is reduced to 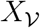 and response matrix is reduced to 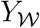. The coefficient matrix is reduced to a smaller and more sparse matrix 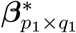 with edge sparsity indicated by the matrix **U**_*P*_1_×*q*_1__, where *U_lh_* = 1 if 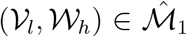 and *U_lh_* = 0 if 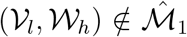 from the screening stage. One can then apply a regularization method to refine the final pool of the most important pairs. In this paper, we consider the following objective function with both *ℓ*_1_ and *ℓ*_2_ norm penalty to estimate 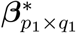:

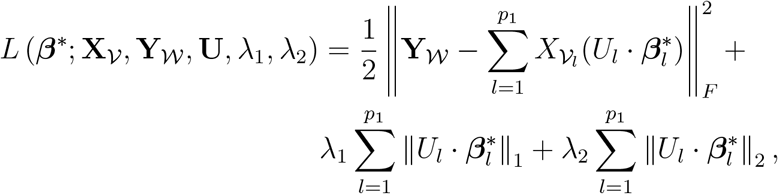

where *U_l_* = (*U*_*l*1_,…, *U*_*lq*1_) is the *l*-th row of 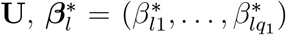 is the *l*-th row of 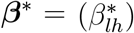; || · ||_*F*_ denotes the Frobenius norm of matrices; || · ||_1_ and || · ||_2_ are the *ℓ*_1_ and *ℓ*_2_ norms for vectors, respectively, and “·” stands for Hadamard product. The *ℓ*_1_ norm penalty on the ***β***\* matrix ensures the edge-wise sparsity of the multivariate regression model while penalizing the *ℓ*_2_ norm of each row of the ***β***\* matrix constrains the total number of predictors kept in the model and achieves node-wise sparsity. Minimizing the above objective function gives us 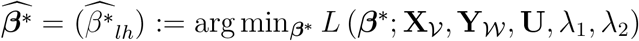, where *λ*_1_ and *λ*_2_ are the tuning parameters. The final estimate of the *p* × *q* coefficient matrix 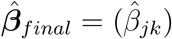 can be obtained as 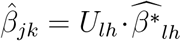 if 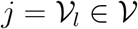 and 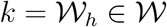, and 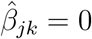 otherwise. It is worth pointing out that the existence of **U** makes sure that only (*X_j_, Y_k_*) pairs that survive after screening step will contribute to the objective function in regularization step. The matrix **U** is usually very sparse which helps speed up the optimization even when *p*_1_ and *q*_1_ are relatively large. We solve the above optimization problem using the R package remMap (Peng et al., 2010). The optimal *λ*_1_ and *λ*_2_ are chosen by 5-fold cross-validation.

#### Algorithm 1 rPCor algorithm

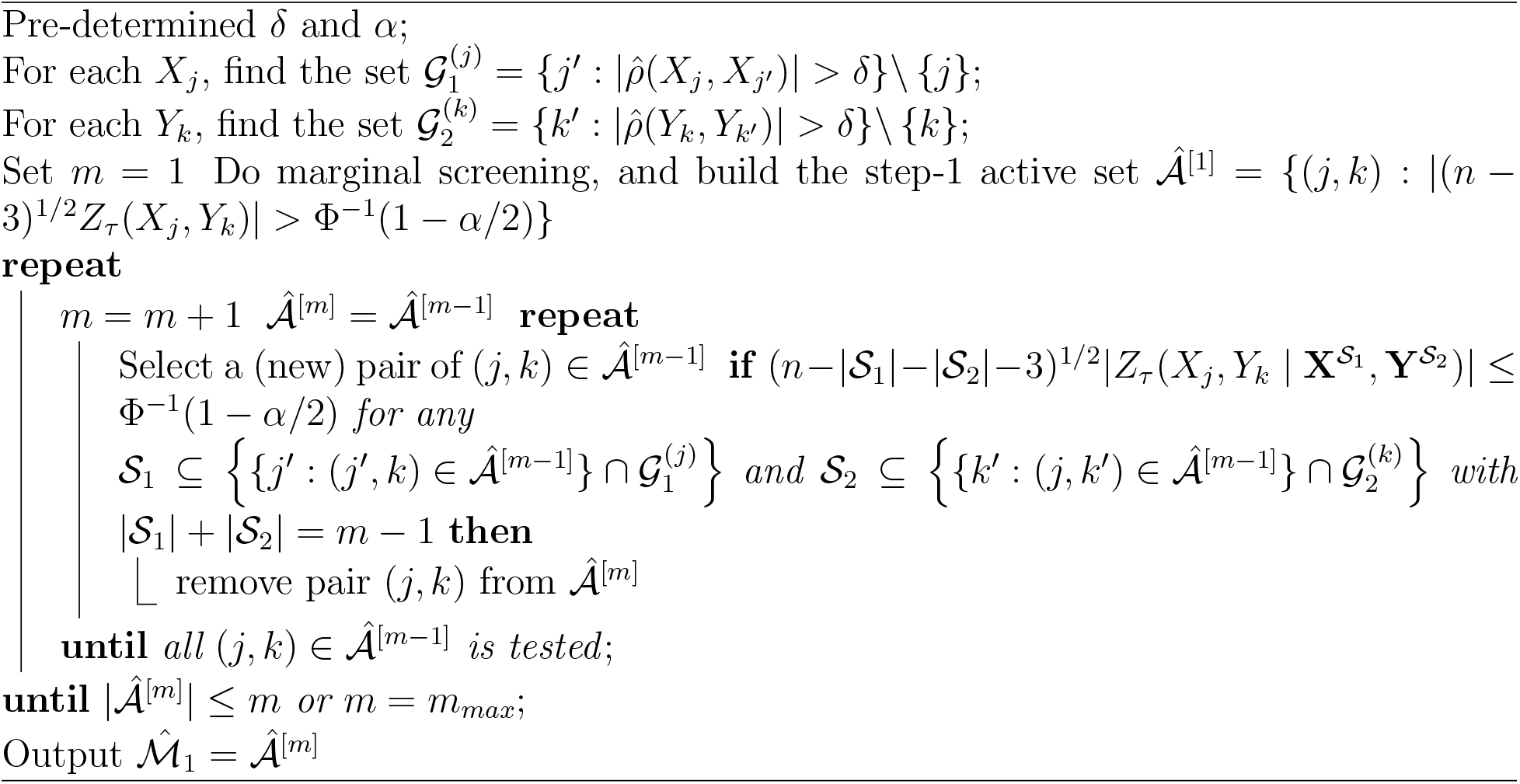

## 3 Simulation

### 3.1 Simulation settings

In this section, we conducted comprehensive simulation studies to show the performance of our screening method. To benchmark our method, we compared rPCor to other popular screening methods in linear models, including SIS (Fan and Lv, 2008), CIS (He et al., 2019) and PC-simple (Bühlmann et al., 2010), as well as a model-free screening method, DC-SIS (Li et al., 2012). For fair comparison, screening methods that use the model size tuning were reformulated to use the threshold tuning by *α*.

#### Scenario I

In this scenario, we evaluated the screening performance of our method under Gaussian assumption. We simulated the epigenetic data from a multivariate normal distribution **X**_*n*×*p*_ ~ *MVN*(0, **Σ**_*X*_), where ∑_*X,jj*_ = 1 and 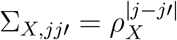 for 1 ≤ *j* ≠ *j*’ ≤ *p*. For ***β***_*p*×*q*_ matrix, we assumed a majority of coefficients are zero (i.e. no regulatory relationship for a majority of epigenetic-gene pairs) and only *s* pairs had nonzero coefficients, the positions of which were randomly distributed in the matrix but restricted to *s_X_* epigenetic features and *s_Y_* genes. Lastly, we simulated the gene expression data by **Y**_*n*×*q*_ ~ *MVN*(**X*β*, Σ_Y_**), where **Σ_*Y*_** is the covariance matrix specified in two ways:

(IA). Toeplitz within modules: we assumed **Σ**_*Y*_ to have block-diagonal structure with non-overlapped gene modules. Each block is a Toeplitz matrix with parameter *ρ_Y_* and the between block correlation is 0.
(IB). Big Toeplitz: we did not assume any gene modules and set *Σ_Y,kk_* = 1 and 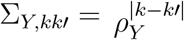 for 1 ≤ *k* ≠ *k*’ ≤ *q*.

In both cases, we assumed *n* = 500, *p* = 5000, *q* = 2000, *s* = 200, *s_X_* = 100, *s_Y_* = 50 and the signal strength of nonzero coefficients *β* = 2. For IA, we assumed 36 independent modules with module size equal to 50 and the rest as scattered genes not belonging to any modules. For inter-feature correlation *ρ_X_* and *ρ_Y_*, we considered different combinations, i.e. (*ρ_X_*, *ρ_Y_*) ∈ {(0.2, 0.2), (0.8,0.2), (0.2, 0.8), (0.8, 0.8)}.

#### Scenario II

In the second scenario, we aimed to assess the robustness of our method when the Gaussian assumption was violated, which is common in the high-dimensional biomedical data. We generated epigenetic predictors **X**_*n*×*p*_ from Pareto distribution with both shape and scale parameters equal to 1. In this example, we assumed *n* = 100, *p* = 1000, *q* = 200, *s* = 9, *s_X_* = 9 and *s_Y_* = 3.

Each of the simulations was replicated for *B* = 100 times. For scenario I, we assessed the true positive rate (or sensitivity) and the empirical false discovery rate at both edge-level (epigenetic-gene pairs) and node-level (epigenetic features or genes). Nodes that have at least one true edge were regarded as true nodes. Unlike the evaluation of regularization methods that looked at both sensitivity and specificity, for screening methods, we assessed whether they can remove as many false positives without losing any signals (i.e. maintain a high sensitivity). For scenario II, we mainly assessed the sensitivity of the methods in the presence of heavy-tailed data.

### 3.2 Simulation results

Figure 3 shows both the edge-wise and node-wise results for Scenario IA when (*ρ_X_, ρ_Y_*) = (0.8, 0.8). As *α* became more stringent, all methods started to remove more edges. Our method rPCor kept almost all the true edges as the other methods did (upper) while significantly reducing the number of false positive edges (lower). The advantage remained as we assessed the methods at node level, where rPCor kept all true nodes but significantly reduced the number of predictors and responses left. Since rPCor considered the between-response correlations, it removed much more false positive responses (Y-nodewise) than the other methods. Although CIS removed even more false positives, it also lost many true positives at the same time, which is unacceptable during the screening stage. As the correlations *ρ_X_* and *ρ_Y_* dropped, the advantage of rPCor became less prominent as expected (Fig S2-S4). Scenario IB showed a similar trend where rPCor had similar sensitivity as the other methods while keeping a much lower false discovery rates (Fig S5).

**Figure 3:**
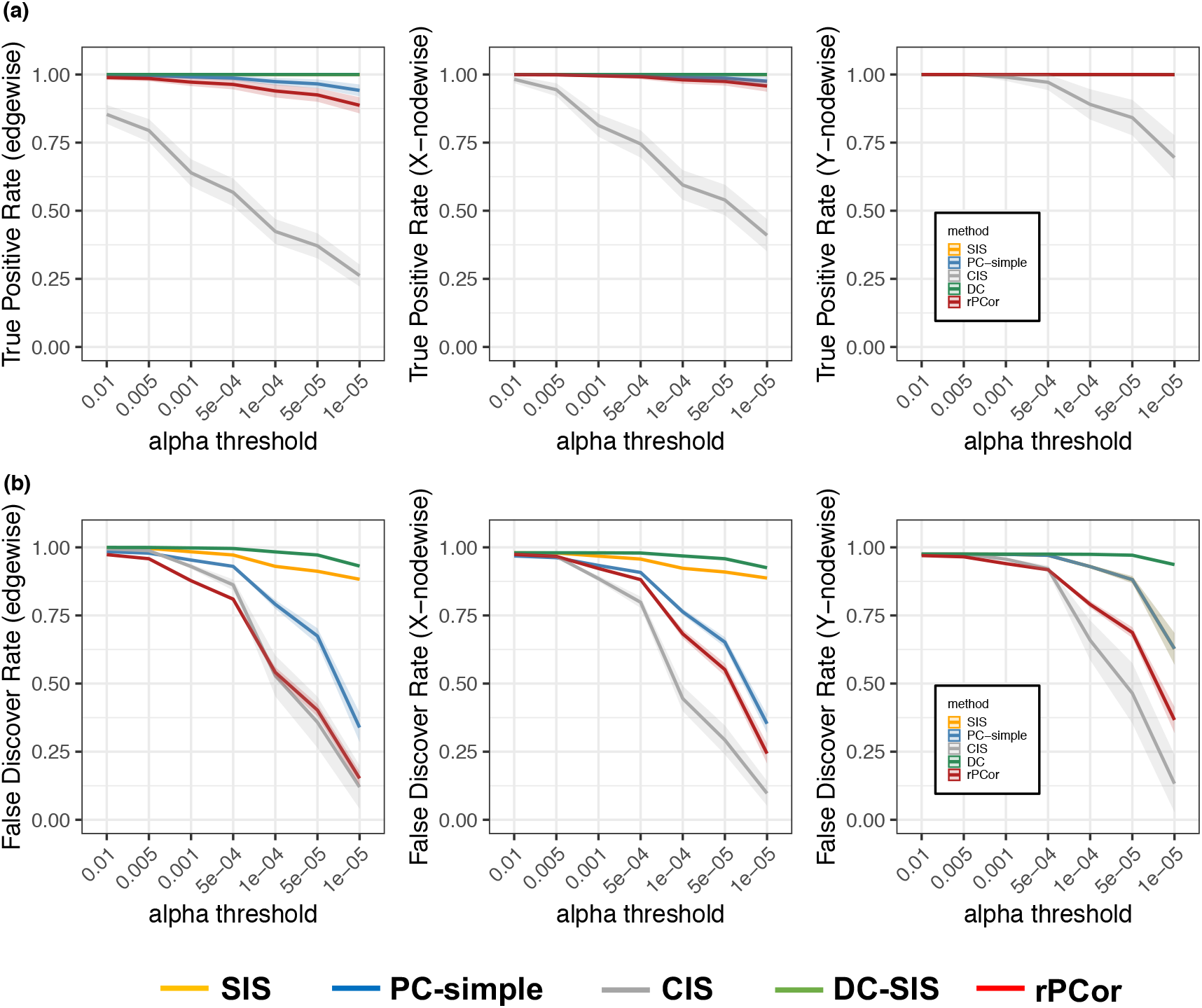
Simulation results for Scenario (IA) with (*ρ_X_, ρ_Y_*) = (0.8,0.8). (a) True positive rates comparison for edgewise, X-nodewise (predictors) and Y-nodewise (responses) results. (b) False discover rates comparison for edgewise, X-nodewise and Y-nodewise results. The mean and confidence band of each metric at different *a* levels are presented here.

In Scenario II, rPCor had much higher sensitivity than the other non-robust screening methods, indicating the necessity of performing robustification on partial correlation in the presence of heavy-tailness (Fig S6). Table S1 summarized the average computational time of all screening methods for different scenarios. Our screening method was computationally much faster than other partial correlation-based methods (CIS and PC-simple) and not any worse than the marginal correlation based screening method SIS. When the sets 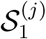 and 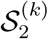 are small, the iterative algorithm had a computational complexity approximately the same as the marginal correlation screening (bounded by *O*(*npq*)).

## 4 Real data

### 4.1 Long non-coding RNA regulation in kidney cancer

We applied our method to the TCGA Pan-kidney cohort (including kidney chromophobe (KICH), kidney renal clear cell carcinoma (KIRC), and kidney renal papillary cell carcinoma (KIRP)) that collects both lncRNA data and gene expression data for kidney cancer patients. LncRNAs are essential regulators of genes in major pathways of cell growth, proliferation, differentiation and survival, and are critical to the tumour formation, progression and pathogenesis of kidney cancer (Zhou et al., 2014; Martens-Uzunova et al., 2014). Our purpose is to identify important lncRNA regulators of genes in kidney cancer over the whole genome.

The lncRNA expression data (in Reads Per Million mapped reads, or RPM) and the gene expression data (in Reads Per Kilobase of transcript per Million mapped reads, or RPKM) of TCGA Pan-kidney cohort, both measured by RNA-sequencing, were downloaded from The Atlas of Noncoding RNAs in Cancer (TANRIC; Li et al. (2015a)) and LinkedOmics (Vasaikar et al., 2018), respectively. We carefully preprocessed the data by matching the samples, filtering out features with low expression while keeping lncRNAs with mean values greater than 0.3 and genes with mean expression values greater than 5 only following the general guideline (Li et al., 2015a; Ricketts et al., 2018). The processed data included *p* = 2170 lncRNAs and *q* = 13408 genes for *n* = 712 kidney cancer patients.

Existing regularization methods cannot afford the computation for such large scale data, so we will perform screening methods to reduce dimension first. We applied rPCor, SIS, DC-SIS and PC-simple to the data. Table 2 summarizes the number of lncRNAs, genes as well as lncRNA-gene pairs left after screening. The model after screening by rPCor is much more sparse than that by other methods, leaving a total of 7528 lncRNA-gene pairs that included *p*_1_ = 1652 lncRNAs and *q*_1_ = 5274 genes. On the other hand, a much larger number of lncRNA-gene pairs and unique lncRNAs and genes remained after screening by the other methods under the same threshold level, so regularization methods can still not be applied.

**Table 2:** Screening and regularization results of different methods for Pan-kidney example method # of pairs left # of lncRNAs left # of genes left

| method | # of pairs left | # of lncRNAs left | # of genes left |
| --- | --- | --- | --- |
| original data | 29095360 | 2170 | 13408 |
| SIS | 9715838 | 2170 | 13406 |
| DC-SIS | 17607189 | 2170 | 13408 |
| PC-simple | 59950 | 1598 | 13406 |
| rPCor | 7528 | 1652 | 5274 |
| rPCor + regularization | 296 | 46 | 273 |

We then applied the proposed regularization method to the set after screening by rPCor and selected 296 lncRNA-gene pairs with 46 unique lncRNAs and 273 unique genes as our final selection results. It was notable that several lncRNAs regulated multiple genes both positively and negatively (see Fig S7-9 for the visualization of identified regulatory networks by using the tool MSLCRN (Zhang et al., 2019)), a pattern common to lncRNA regulation (Jiang et al., 2018). Many of the identified lncRNAs and genes were significantly associated with different subtypes, pathological stages and survival time of kidney caner, and can potentially serve as candidate biomarkers for clinical prognosis (Table S5, Fig S10-11). We then performed pathway enrichment analyses for the 273 genes selected using four pathway databases: GO (Ashburner et al., 2000), KEGG (Kanehisa and Goto, 2000), BioCarta (Nishimura, 2001) and Reactome (Fabregat et al., 2017). Among the top 10 enriched pathways sorted by their corresponding -log10(p-value) from the Fisher’s exact test (Fig 4), we found pathways of important biological processes critical to renal cell carcinoma such as Reactome Integrin cell surface interactions and KEGG Natural killer cell mediated cytotoxicity (Akçay, 2021; Li et al., 2019), showing the advantage of our method in identifying critical disease related gene markers regulated by lncRNAs.

**Figure 4:**
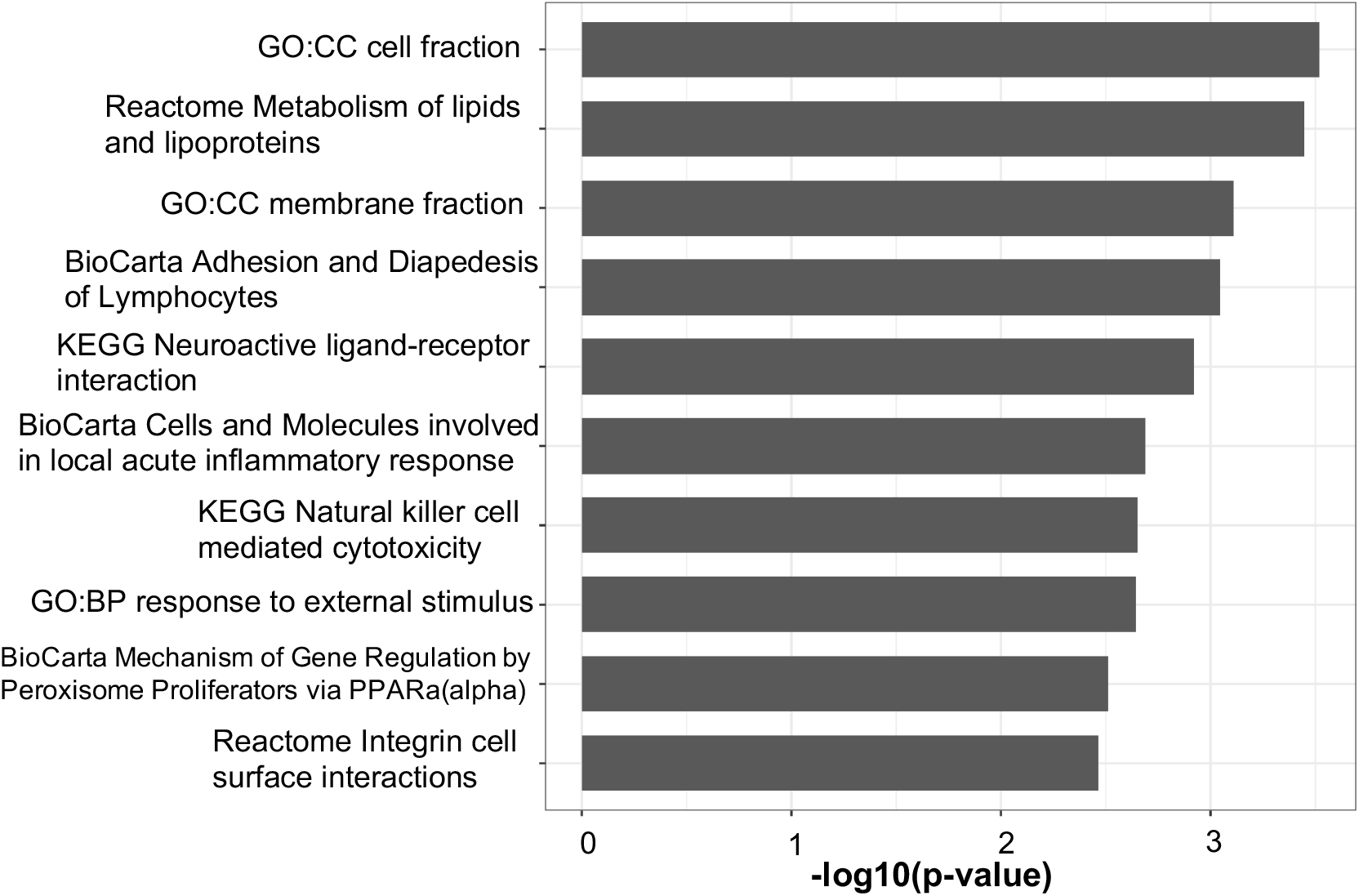
Top 10 pathways enriched with the 273 genes finally selected in kidney cancer lncRNA-gene regulation example.

### 4.2 DNA methylation regulation in Glioblastoma Multiforme

In a second example, we applied our method to study DNA methylation regulation of genes in Glioblastoma Multiforme (GBM) using data from TCGA. GBM is one of the most aggressive types of malignant primary brain tumor, identifying the DNA methylation features (e.g. methylated CpG sites) that regulate gene expression in GBM helps understand the disease mechanism and find potential therapeutic targets. Detailed data preprocessing and comparison results can be found in Supplement section 8. “rPCor+regularization” selected a final pool of 481 CpG-gene pairs that included 180 unique CpG sites and 429 unique genes (Table S2). Several identified genes and CpG sites were significantly associated with the survival time of GBM implying their potential roles as prognostic markers (Table S6, Fig S12). We found an enrichment of transcription factors (TFs) among the 429 genes selected (Table S3: *N* = 42, Fisher’s Exact Test *p* < 0.05). Both cis-regulation (TFs regulated by methylated CpG sites in promoter region) and trans-regulation (TFs regulated by CpG sites out of promoter region) were identified (Table S4) and the associations were mainly negative among the top associated pairs (Table S7) implying the down-regulatory roles DNA methylation played on gene activity. DNA methylation regulates gene expression by recruiting gene repressor proteins or inhibiting the binding of TFs (Moore et al., 2013). The role of TFs as readers and effectors of DNA methylation has been well documented in literature, where DNA methylation interacts with TFs which in turn regulate the activity of many other genes (Zhu et al., 2016).

## 5 Discussion

In this paper, we proposed a robust partial correlation based screening method to detect epigenetic regulators of gene expression over the whole genome. Our method includes data-driven procedures to determine the conditional sets in partial correlation and the optimal screening threshold, and has robust screening performance in the presence of heavy-tailed data. Existing screening methods target at univariate or low-dimensional responses and perform screening only at node-level. Our screening method reduces the high dimensionality of both predictor and response and manages to screen out noises at both node and edge levels, which work perfectly for epigenetic-gene regulation problem. We have shown the advantage of our method in both screening performance and computation via simulations and real data application to understand the regulatory roles of lncRNA and DNA methylation on essential genes in cancer studies.

Most epigenetic studies to date are on a small scale and restricted to cis epigenetic regulators (Allis and Jenuwein, 2016). With recent development of high-throughput technology, more and more epigenome-wide studies are conducted producing a large amount of epigenetic and gene expression data in public domains. However, there lacks computationally efficient and statistically rigorous methods that jointly analyze these data to investigate their regulatory relationships which will deepen our understanding of the molecular mechanism of disease. Our method is the first screening method that targets at this big data integration problem and allows users to quickly narrow down the genome-wide analysis to a reduced set of epigenetic and gene markers critical to the underlying biology of the disease for selection in the next stage. Our method is generalizable to a wide variety of similar big data integration problems, including but not limited to imaging-genetics, eQTL and multi-omics QTL, etc. The actual screening performance will need to be evaluated case by case in future studies.

Since Fan and Lv (2008) proposed the first sure screening method, numerous screening methods using different statistics and targeting at different models have been developed. However, none of them have ever targeted at a multivariate regression model with highdimensional predictors and responses and considered the screening of both predictors and responses. We proposed the first screening method of this kind within a linear model. The method is readily extensible to other non-linear models or be model-free as in many other extensions of univariate screening methods, which can potentially overcome many existing issues of modern biomedical data. In addition, future research will focus on building more complete theoretical basis of the method and its extension.

## Acknowledgements

Research reported in this publication was supported by the National Institute on Drug Abuse (NIDA) of National Institute of Health under the award number 1DP1DA048968-01 to S.C. and T.M., by the University of Maryland MPower Brain Health and Human Performance seed grant to H.K., S.C. and T.M.

## Notes

https://github.com/kehongjie/rPCor

### Competing Interest Statement

The authors have declared no competing interest.

## References

Aguet, F. and Muñoz Aguirre, M. (2017). Genetic effects on gene expression across human tissues. Nature, 550:204–213.

Akçay, S. (2021). Integrated network analysis of the potential molecular biomarkers and key pathways in clear renal cell carcinoma (ccrcc). Journal of Applied Biological Sciences, 15(3):342–351.

Allis, C. D. and Jenuwein, T. (2016). The molecular hallmarks of epigenetic control. Nature Reviews Genetics, 17(8):487–500.

Ashburner, M., Ball, C. A., Blake, J. A., Botstein, D., Butler, H., Cherry, J. M., Davis, A. P., Dolinski, K., Dwight, S. S., Eppig, J. T., et al. (2000). Gene ontology: tool for the unification of biology. Nature genetics, 25(1):25–29.

Baylin, S. B. and Jones, P. A. (2016). Epigenetic determinants of cancer. Cold Spring Harbor perspectives in biology, 8(9):a019505.

Bühlmann, P., Kalisch, M., and Maathuis, M. H. (2010). Variable selection in highdimensional linear models: partially faithful distributions and the pc-simple algorithm. Biometrika, 97(2):261–278.

Cheng, Y., He, C., Wang, M., Ma, X., Mo, F., Yang, S., Han, J., and Wei, X. (2019). Targeting epigenetic regulators for cancer therapy: mechanisms and advances in clinical trials. Signal transduction and targeted therapy, 4(1):1–39.

Di He, Y. Z. and Zou, H. (2021). On sure screening with multiple responses. Statistica Sinica, 31:1749–1777.

Fabregat, A., Sidiropoulos, K., Viteri, G., Forner, O., Marin-Garcia, P., Arnau, V., D’Eustachio, P., Stein, L., and Hermjakob, H. (2017). Reactome pathway analysis: a high-performance in-memory approach. BMC bioinformatics, 18(1):1–9.

Fan, J. and Lv, J. (2008). Sure independence screening for ultrahigh dimensional feature space. Journal of the Royal Statistical Society: Series B (Statistical Methodology), 70(5):849–911.

Fan, J. and Song, R. (2010). Sure independence screening in generalized linear models with np-dimensionality. The Annals of Statistics, 38(6):3567–3604.

Gibney, E. and Nolan, C. (2010). Epigenetics and gene expression. Heredity, 105(1):4–13.

He, K., Kang, J., Hong, H. G., Zhu, J., Li, Y., Lin, H., Xu, H., and Li, Y. (2019). Covariance-insured screening. Computational statistics & data analysis, 132:100–114.

Jiang, W., Zhan, H., Jiao, Y., Li, S., and Gao, W. (2018). A novel lncrna-mirna-mrna network analysis identified the hub lncrna rp11-159f24. 1 in the pathogenesis of papillary thyroid cancer. Cancer medicine, 7(12):6290–6298.

Kanehisa, M. and Goto, S. (2000). Kegg: kyoto encyclopedia of genes and genomes. Nucleic acids research, 28(1):27–30.

Ke, Y., Minsker, S., Ren, Z., Sun, Q., and Zhou, W.-X. (2019). User-friendly covariance estimation for heavy-tailed distributions. Statistical Science, 34(3):454–471.

Li, F., Guo, P., Dong, K., Guo, P., Wang, H., and Lv, X. (2019). Identification of key biomarkers and potential molecular mechanisms in renal cell carcinoma by bioinformatics analysis. Journal of Computational Biology, 26(11):1278–1295.

Li, J., Han, L., Roebuck, P., Diao, L., Liu, L., Yuan, Y., Weinstein, J. N., and Liang, H. (2015a). Tanric: an interactive open platform to explore the function of lncrnas in cancer. Cancer research, 75(18):3728–3737.

Li, R., Zhong, W., and Zhu, L. (2012). Feature screening via distance correlation learning. Journal of the American Statistical Association, 107(499):1129–1139.

Li, Y., Nan, B., and Zhu, J. (2015b). Multivariate sparse group lasso for the multivariate multiple linear regression with an arbitrary group structure. Biometrics, 71(2):354–363.

Liu, J., Zhong, W., and Li, R. (2015). A selective overview of feature screening for ultrahigh-dimensional data. Science China Mathematics, 58(10):1–22.

Ma, T., Ren, Z., and Tseng, G. C. (2020). Variable screening with multiple studies. Statistica Sinica, 30(2):925–953.

Martens-Uzunova, E. S., Böttcher, R., Croce, C. M., Jenster, G., Visakorpi, T., and Calin, G. A. (2014). Long noncoding rna in prostate, bladder, and kidney cancer. European urology, 65(6):1140–1151.

Meinshausen, N. and Bühlmann, P. (2010). Stability selection. Journal of the Royal Statistical Society: Series B (Statistical Methodology), 72(4):417–473.

Moore, L. D., Le, T., and Fan, G. (2013). Dna methylation and its basic function. Neuropsychopharmacology, 38(1):23–38.

Nishimura, D. (2001). Biocarta. Biotech Software & Internet Report: The Computer Software Journal for Scient, 2(3):117–120.

Peng, J., Wang, P., Zhou, N., and Zhu, J. (2009). Partial correlation estimation by joint sparse regression models. Journal of the American Statistical Association, 104(486):735–746.

Peng, J., Zhu, J., Bergamaschi, A., Han, W., Noh, D.-Y., Pollack, J. R., and Wang, P. (2010). Regularized multivariate regression for identifying master predictors with application to integrative genomics study of breast cancer. The annals of applied statistics, 4(1):53.

Ricketts, C. J., De Cubas, A. A., Fan, H., Smith, C. C., Lang, M., Reznik, E., Bowlby, R., Gibb, E. A., Akbani, R., Beroukhim, R., et al. (2018). The cancer genome atlas comprehensive molecular characterization of renal cell carcinoma. Cell reports, 23(1):313–326.

Vasaikar, S. V., Straub, P., Wang, J., and Zhang, B. (2018). Linkedomics: analyzing multi-omics data within and across 32 cancer types. Nucleic acids research, 46(D1):D956–D963.

Zhang, J., Le, T. D., Liu, L., and Li, J. (2019). Inferring and analyzing module-specific lncrna-mrna causal regulatory networks in human cancer. Briefings in bioinformatics, 20(4):1403–1419.

Zhou, S., Wang, J., and Zhang, Z. (2014). An emerging understanding of long noncoding rnas in kidney cancer. Journal of cancer research and clinical oncology, 140(12):1989–1995.

Zhu, H., Wang, G., and Qian, J. (2016). Transcription factors as readers and effectors of dna methylation. Nature Reviews Genetics, 17(9):551–565.

Zhu, L.-P., Li, L., Li, R., and Zhu, L.-X. (2011). Model-free feature screening for ultrahigh-dimensional data. Journal of the American Statistical Association, 106(496):1464–1475.

