## Supplement for "High-dimension to high-dimension screening for detecting genome-wide epigenetic regulators of gene expression"

January 2022

### 1 Review of popular screening methods

In this section, we will review some of the existing popular screening methods for high dimensional data, that are more related to our method. All of the methods consider a univariate response except DC-SIS can be used for low-dimensional multivariate response.

#### 1.1 SIS

Sure Independence Screening (SIS) (Fan and Lv, 2008) is the first variable screening method ever proposed and applies to a linear model. The idea of SIS is simple and straightforward: selecting predictors using their marginal sample correlations with the response, i.e. $\rho(Y, X_j), j \in \{1, ..., p\}$. To identify the nonzero coefficients in vector $\beta$, SIS sorts the absolute values of $p$ marginal correlations $\rho(Y, X_j)$ and define the sub-model as

$$\widehat{\mathcal{M}_{SIS}} = \{1 \leq j \leq p : |\rho(Y, X_j)| \text{ is among the first } [\gamma n] \text{ largest of all } \},$$

where $\gamma \in (0, 1)$ is pre-determined and $[\gamma n]$ denotes the integer part of $\gamma n$. With SIS, the model size after screening is $d = [\gamma n]$, and Fan and Lv (2008) recommends using a theoretically guided $d = n - 1$ or $d = n/\log n$ depending on the order of sample size $n$.

#### 1.2 DC-SIS

The sure independence screening based on the distance correlation (DC-SIS) (Li et al., 2012) extends SIS to a more general setting and does not require model specification for responses or predictors. Based on Székely et al. (2007), a natural estimator of sample distance covariance is

$$\widehat{d\,\mathrm{cov}}^2(\mathbf{x}, \mathbf{y}) = \hat{S}_1 + \hat{S}_2 - 2\hat{S}_3,$$

where

$$\hat{S}_1 = \frac{1}{n^2} \sum_{k=1}^{n} \sum_{l=1}^{n} \|\mathbf{x}_k - \mathbf{x}_l\|_{d_x} \|\mathbf{y}_k - \mathbf{y}_l\|_{d_y},$$

$$\hat{S}_2 = \frac{1}{n^2}\sum_{k=1}^{n}\sum_{l=1}^{n}\|\mathbf{x}_k - \mathbf{x}_l\|_{d_x}\frac{1}{n^2}\sum_{k=1}^{n}\sum_{l=1}^{n}\|\mathbf{y}_k - \mathbf{y}_l\|_{d_y} \text{ and}$$

$$\hat{S}_3 = \frac{1}{n^3}\sum_{k=1}^{n}\sum_{l=1}^{n}\sum_{m=1}^{n}\|\mathbf{x}_k - \mathbf{x}_l\|_{d_x}\|\mathbf{y}_m - \mathbf{y}_l\|_{d_y}.$$

Then the sample distance correlation can be defined as

$$\widehat{dc}(\mathbf{x}, \mathbf{y}) = \frac{\widehat{d\,\mathrm{cov}}(\mathbf{x}, \mathbf{y})}{\sqrt{\widehat{d\,\mathrm{cov}}(\mathbf{x}, \mathbf{x})\widehat{d\,\mathrm{cov}}(\mathbf{y}, \mathbf{y})}}.$$

Similar to SIS, DC-SIS selects a set of important predictors with large values of $\widehat{dc}(X_j, Y)$. That is, DC-SIS will select the sub-model as

$$\widehat{\mathcal{M}_{DC}} = \left\{j : \widehat{dc}^2(X_j, Y) \geq cn^{-\kappa}, \text{ for } 1 \leq j \leq p\right\},$$

where $c$ and $\kappa$ are pre-specified threshold guided by the theory.

It is worth mentioning that under normality assumption, using DC-SIS for screening is equivalent to using SIS as DC is monotonically increasing in Pearson correlation in this case; and DC-SIS is more effective than SIS when there exist non-linear relationship between two random variables. However, the computation burden of DC is extremely heavy while calculating Pearson correlation is simple and fast.

#### 1.3 PC-simple

Bühlmann et al. (2010) proposes a partial correlation based algorithm (PC-simple) that iteratively screen out predictors that has small partial correlation $\rho_(Y, X_j|X^{(\mathcal{S})})$ with the response. We first define the Fisher's Z-transformation for sample partial correlation as

$$Z(Y, X_j \mid X^{(\mathcal{S})}) = \frac{1}{2}\log\left\{\frac{1 + \hat{\rho}(Y, X_j \mid X^{(\mathcal{S})})}{1 - \hat{\rho}(Y, X_j \mid X^{(\mathcal{S})})}\right\}.$$

PC-simple starts with building the active set at marginal level or zero-order, i.e.

$$\hat{\mathcal{A}}^{[1]} = \{j : |(n-3)^{1/2}Z(X_j, Y_k)| > \Phi^{-1}(1 - \alpha/2)\},$$

where $\Phi(\cdot)$ is the standard Normal cumulative distribution function, $\alpha$ is the threshold chosen to be the unadjusted significance level commonly used, e.g. $\alpha = 0.05$. Then the order of partial correlation keeps increasing and the active set will be updated as

$$\begin{aligned}\hat{\mathcal{A}}^{[m]} = \{j \in \hat{\mathcal{A}}^{[m-1]} : (n - |\mathcal{S}| - 3)^{1/2}|Z(Y, X_j \mid X^{(\mathcal{S})}| > \Phi^{-1}(1 - \alpha/2) \\ \text{for all } \mathcal{S} \subseteq \mathcal{A}^{[m-1]}\backslash\{j\} \text{ with } |\mathcal{S}| = m - 1\}.\end{aligned}$$

The maximum value of $m$ that is reached in this algorithm is called $m_{reach}$ and can be written as $m_{\text{reach}} = \min\left\{m : \left|\mathcal{A}^{[m]}\right| \leqslant m\right\}$. Thus, the sub-model that PC-simple selects is simply

$$\widehat{\mathcal{M}_{PC}} = \hat{\mathcal{A}}^{[m_{reach}]}.$$

#### 1.4 CIS

To leverage inter-predictor dependence while maintaining computational feasibility, He et al. (2019) proposes Covariance Insured Screening (CIS) which utilizes semi-partial correlation and restricts the conditional set within some pre-determined blocks. It starts with identify $G$ disconnected blocks ($\hat{\mathcal{S}}_g, 1 \leq g \leq G$) in predictors by thresholding the sample covariance matrix. For each $j \in \hat{\mathcal{S}}_g$, one can find the projection matrix onto the space spanned by $\mathbf{x}_{\hat{\mathcal{S}}_g}$ as

$$\Pi_{\hat{\mathcal{S}}_g\backslash\{j\}} = \mathbf{x}_{\hat{\mathcal{S}}_g\backslash\{j\}} \left(\mathbf{x}^T_{\hat{\mathcal{S}}_g\backslash\{j\}}\mathbf{x}_{\hat{\mathcal{S}}_g\backslash\{j\}}\right)^{-1} \mathbf{x}_{\hat{\mathcal{S}}_g\backslash\{j\}}.$$

Then the block-wise sample semi-partial correlation can be calculated as

$$\hat{\rho^*}\left(Y_i, X_{i,j} \mid \mathbf{X}_{i,\hat{\mathcal{S}}_g\backslash\{j\}}\right) = \frac{\mathbf{x}_j^T\left(\mathbf{I}_n - \Pi_{\hat{\mathcal{S}}_g\backslash\langle\rangle}\right)\mathbf{y}}{\sqrt{\mathbf{x}_j^T\left(\mathbf{I}_n - \Pi_{\hat{\mathcal{S}}_g\backslash\{j\}}\right)\mathbf{x}_j\sqrt{\mathbf{y}^T\mathbf{y}}}}.$$

Similarly, CIS will then sort all predictors based on their absolute values of semi-partial correlations and identify the submodel as

$$\widehat{\mathcal{M}_{CIS}} = \left\{j \in \hat{\mathcal{S}}_g, 1 \leq g \leq G : \left|\hat{\rho}\left(Y_i, X_{i,j} \mid \mathbf{X}_{i,\hat{\mathcal{S}}_g\backslash\{j\}}\right)\right| > v\right\},$$

where $v$ is a pre-defined threshold guided by the theory.

### 2 Paritial faithfulness assumption and connection between partial correlation and regression coefficient

Under a standard univariate linear model:

$$y = \sum_{j=1}^{p} X_j\beta_j + \epsilon,\ \epsilon \sim N(0, \sigma^2),$$

where $\beta_j$ is the regression coefficient for $j$th variable $X_j$. Bühlmann et al. (2010) first defined the partial faithfulness assumption for the above linear model if for every $j \in \{1, 2, \ldots, p\}$:

$$\rho(y, X_j|X_S) = 0 \text{ for some } S \in \{j\}^C \to \beta_j = 0.$$

They showed that if a linear model satisfied: (1) $cov(X) = \Sigma_X$ is positive definite; (2) the nonzero regression coefficients $\{\beta_j : \beta_j \neq 0\} \sim f(b)db$, then the partial faithfulness holds almost surely. This condition justifies the rationale of using partial correlation as a screening statistics to identify the active set of variables with nonzero coefficients.

Similar connection between partial correlation and regression coefficient have also been shown in Witten and Tibshirani (2009), where they linked the problem of regressing $y$ onto $X$ to the problem of estimating inverse covariance matrix (or precision matrix) of $y$ and $X$ in Gaussian Graphical Model (GGM). Suppose $T = (X\ y)$ and $cov(T) = \Sigma$ and $\Theta = \Sigma^{-1}$ assuming $\Sigma$ is invertible. Let $\Theta = \begin{pmatrix} \Theta_{XX} & \Theta_{Xy} \\ \Theta_{Xy}^T & \Theta_{yy} \end{pmatrix}$, it can be shown that $\beta = -\Theta_{Xy}/\Theta_{yy}$, where $\Theta_{Xy}$ is the off-diagonal portion of $\Theta$. Since the partial correlation $\rho(X_j, y|X^{\{j\}^C}) = -\frac{\Theta_{X_j y}}{\Theta_{X_j X_j}\Theta_{yy}}$, we have $\rho(X_j, y|X^{\{j\}^C}) = 0 \leftrightarrow \Theta_{X_j y} = 0 \leftrightarrow \beta_j = 0$. This is a special case of the partial faithfulness assumption when $S = \{j\}^C$.

For a multivariate regression model:

$$Y_{n\times q} = X_{n\times p}\beta_{p\times q} + E_{n\times q}, E \sim N(0, \Sigma_Y \otimes I_n),$$

where $\beta$ is the $p \times q$ matrix of regression coefficients, $cov(X) = \Sigma_X$, $\Sigma_Y$ is the $q \times q$ covariance matrix of $Y$. Suppose $T = (X\ Y)$ and $cov(T) = \Sigma$, with block-wise decomposition, $\Sigma = \begin{pmatrix} \Sigma_X & \Sigma_{XY} \\ \Sigma_{YX} & \Sigma_Y \end{pmatrix}$. Assuming $\Sigma$ is strictly positive definite thus invertible, and let $\Theta = \Sigma^{-1} = \begin{pmatrix} \Theta_X & \Theta_{XY} \\ \Theta_{YX} & \Theta_Y \end{pmatrix}$. It can be shown that the above multivariate regression model can be reparameterized in the form of a conditional Gaussian Graphical Model (cGGM) (Sohn and Kim, 2012; Chiquet et al., 2017):

$$Y|X \sim N(-X\Theta_{XY}\Theta_Y^{-1}, \Theta_Y^{-1} \otimes I_n),$$

where $\beta = -\Theta_{XY}\Theta_Y^{-1}$. The partial correlation $\rho(X_j, Y_k|X^{\{j\}^C}, Y^{\{k\}^C}) = -\frac{\Theta_{X_j Y_k}}{\Theta_{X_j X_j}\Theta_{Y_k Y_k}}$, so we have $\rho(X_j, Y_k|X^{\{j\}^C}, Y^{\{k\}^C}) = 0 \leftrightarrow \Theta_{X_j Y_k} = 0$, thus there is a one-to-one connection between the regression coefficient and the partial correlation.

#### 3 Tuning of robustification parameter $\tau$

Following Ke et al. (2019) and Wang et al. (2020), we propose a data-adaptive procedure to automatically tune the robustification parameter $\tau$ for each of $X_j$ and $Y_k$ adjusting for the other variables in the conditional sets. Suppose we want to calculate the robust partial correlation $\hat{\rho}_\tau(X_j, Y_k|\mathbf{X}^{\mathcal{S}_1^{(j)}}, \mathbf{Y}^{\mathcal{S}_2^{(k)}})$, we first combine the data column-by-column, i.e. $\mathbf{W} = (X_j, Y_k, \mathbf{X}^{\mathcal{S}_1^{(j)}}, \mathbf{Y}^{\mathcal{S}_2^{(k)}})$ as a $n \times p_0$ matrix where $p_0 = |\mathcal{S}_1^{(j)}| + |\mathcal{S}_2^{(k)}| + 2$. Then we define the pairwise difference by $\{\mathbf{V}_1, \ldots, \mathbf{V}_N\} = \{\mathbf{W}_1 - \mathbf{W}_2, \ldots, \mathbf{W}_{(n-1)} - \mathbf{W}_n\}$, where $\mathbf{W}_i$ is the i-th row of $\mathbf{W}$ and $N = \binom{n}{2}$. Next, for any two columns $(g, h)$ in $\mathbf{W}$, we also define $\left\{Z_{1(gh)} \ldots, Z_{N(gh)}\right\} = \{V_{1g}V_{1h}/2, \ldots, V_{Ng}V_{Nh}/2\}$ such that their covariance $\sigma_{gh} = \mathbb{E}Z_{1(gh)}$. Then a robust estimate of this covariance can be written as

$$\widehat{\sigma}_{gh}^{\tau} = (1/N)\sum_{i=1}^{N} \psi_{\tau_{gh}}\left(Z_{i(gh)}\right),$$

where $\psi_\tau(x) = (|x| \wedge \tau)sign(x)$. It can be shown that an "ideal" choice of $\tau_{gh}$ can be solved from the following equation:

$$\frac{1}{N}\sum_{i=1}^{N} \frac{(Z_{i(gh)}^2 \wedge \tau_{gh}^2)}{\tau_{gh}^2} = \frac{t}{n},$$

where $t = C \log p_0$ as guided by the theory. The choice of $C$ is the same for all pairs $(g, h)$ and can be determined empirically depending on the extent of tail heaviness of the data. When the data is not too skewed (e.g. having small kurtosis), a larger $C$ can be used to choose a small $\tau$, so minimal truncation is implemented to save computational time without impacting the screening performance.

From the above, we can get an optimal estimate of $\tau_{gh}$ for each pair $(g, h)$, respectively. After getting the optimal $\hat{\tau}_{gh}$, we can implement the truncation operator $\psi_\tau(.)$ and obtain a robust estimate of covariance matrix $\hat{\mathbf{\Sigma}}_{\mathbf{W}}$ as well as its precision matrix $\hat{\mathbf{P}}_{\mathbf{W}} = (P_{st})$. Finally, the robust partial correlation can be found as $\hat{\rho}_\tau(X_j, Y_k|\mathbf{X}^{\mathcal{S}_1^{(j)}}, \mathbf{Y}^{\mathcal{S}_2^{(k)}}) = -P_{12}/\sqrt{P_{11}P_{22}}$.

### 4 Alternative method to determine the conditional sets based on unconnected gene modules

Module structure naturally exists in gene expression data, so we can first partition all genes into unconnected modules and restrict the conditional set for each gene to only including genes within the same module, i.e. $\mathcal{G}_2^{(k)} = \widehat{\mathcal{S}_2^{(k)}} = \{k' : Y_{k'} \in \Omega_l\} \setminus \{k\}$ where $Y_k \in \Omega_l$, the $l$th gene module. For gene module detection, correlation based partition procedure along the same lines of the breadth-first search algorithm in graph theory (Even, 2011; He et al., 2019) can be used. Similarly, Witten et al. (2011) and Mazumder and Hastie (2012) discussed finding connected components and identifying blocks in the graphical lasso solution, which also applies to our problem here. Alternatively, more biology-driven algorithms such as weighted gene co-expression network analysis (WGCNA) (Langfelder and Horvath, 2008) and other related methods for co-expression network detection (Van Dam et al., 2018) can also be used. One additional benefit of the module-based approach is that the identified gene module structure might provide new biological interpretation to the regulation problem, e.g. epigenetic features target only within the scope of certain gene modules.

Based on the conditional sets of responses determined from gene modules, we proposed an alternative iterative algorithm "rPCor-module" as an addition to the main algorithm as below:

---

**Algorithm 1:** rPCor-module algorithm

---

Partition $Y's$ into $L$ independent modules $\Omega_1, \Omega_2, \ldots, \Omega_L$;
Pre-determined $\delta$ and $\alpha$;
For each $X_j$, define $\mathcal{G}_1^{(j)} = \{j' : |\hat{\rho}(X_j, X_{j'})| > \delta\} \setminus \{j\}$;
For each $Y_k \in \Omega_l$, define $\mathcal{G}_2^{(k)} = \{k' : Y_{k'} \in \Omega_l\} \setminus \{k\}$ ;

Set $m = 1$;
Do marginal screening, and build the step-1 active set
  $\hat{\mathcal{A}}^{[1]} = \{(j,k) : |(n-3)^{1/2} Z_\tau(X_j, Y_k)| > \Phi^{-1}(1-\alpha/2)\}$;
**repeat**
  $m = m + 1$;
  $\hat{\mathcal{A}}^{[m]} = \hat{\mathcal{A}}^{[m-1]}$;
  **repeat**
    Select a (new) pair of $(j,k) \in \hat{\mathcal{A}}^{[m-1]}$;
    **if** $(n - |\mathcal{S}_1| - |\mathcal{S}_2| - 3)^{1/2} |Z_\tau(X_j, Y_k \mid \mathbf{X}^{\mathcal{S}_1}, \mathbf{Y}^{\mathcal{S}_2})| \le \Phi^{-1}(1-\alpha/2)$ *for any* $\mathcal{S}_1 \subseteq \left\{\{j' : (j',k) \in \hat{\mathcal{A}}^{[m-1]}\} \cap \mathcal{G}_1^{(j)}\right\}$ *and* $\mathcal{S}_2 \subseteq \left\{\{k' : (j,k') \in \hat{\mathcal{A}}^{[m-1]}\} \cap \mathcal{G}_2^{(k)}\right\}$ *with* $|\mathcal{S}_1| + |\mathcal{S}_2| = m - 1$ **then**
      remove pair $(j,k)$ from $\hat{\mathcal{A}}^{[m]}$
  **until** *all* $(j,k) \in \hat{\mathcal{A}}^{[m-1]}$ *is tested*;
**until** $|\hat{\mathcal{A}}^{[m]}| \le m$ *or* $m = m_{max}$;
Output $\hat{\mathcal{M}}_1 = \hat{\mathcal{A}}^{[m]}$

---

### 5 Stability selection procedure to determine the optimal screening threshold $\alpha$

To determine the optimal choice of $\alpha$, we use the idea of Stability Selection that was introduced by Meinshausen and Bühlmann (2010) and proposed a similar procedure as in He et al. (2016). See the algorithm below for details.

---

**Algorithm 2:** To determine optimal threshold $\alpha$.

---

Start with a relatively large $\alpha$.

**Step 1.** Draw $S$ random subsamples of $1, ...n$ of size $[n/2]$ without replacement, run rPCor on each subsample, and compute the selection frequency $\hat{\Pi}_{(j,k)} = \frac{1}{S}\sum_{s=1}^{S} I\left((j,k) \in \hat{\Omega}^{(s)}\right)$, where $\hat{\Omega}^{(s)} = \left\{(j,k) : \hat{\beta}_{jk}^{(s)} \neq 0\right\}$ is the set of selected predictor-response pairs from the s-th subsample and $I$ is the indicator function.

**Step 2.** Randomly permute the response variables $D$ times, and for each permuted sample, we repeat the aforementioned stability calculation and compute $\tilde{\Pi}^{d}_{(j,k)} = \frac{1}{S}\sum_{s=1}^{S} I\left((j,k) \in \tilde{\Omega}^{(d,s)}\right)$, where $\tilde{\Omega}^{(d,s)}$ is the set of selected pairs at d-th permutation and s-th subsample.

**Step 3.** Order the values of $\hat{\Pi}_{(j,k)}$, denote the $\phi$-th ($\phi = 1, ..., pq$) largest value as $\hat{\Pi}_{(\phi)}$, and compute the empirical Bayes false discovery rate for $\hat{\Pi}_{(\phi)}$ (Efron, 2012) as

$$Fdr_{(\phi)} = \min\left\{\frac{1}{D}\frac{\sum_{d=1}^{D}\sum_{j'=1}^{p}\sum_{k'=1}^{q} I\left(\tilde{\Pi}^{d}_{(j',k')} \geq \hat{\Pi}_{(\phi)}\right)}{\sum_{j'=1}^{p}\sum_{k'=1}^{q} I\left(\hat{\Pi}_{(j',k')} \geq \hat{\Pi}_{(\phi)}\right)}, 1\right\}.$$

**Step 4.** Calculate the threshold for selection frequency given a pre-determined value $r \in (0,1)$ as

$$\hat{\Pi}_{thres}(r) = \min\left\{\hat{\Pi}_{(\phi)} : Fdr_{(\phi)} \leq r\right\}.$$

**Step 5.** For the top $\eta = \min\{p, q\}$ predictor-response pairs with largest marginal correlation, check if their minimum selection frequency (for robustness, we focus on those pairs above "median-IQR/2" and exclude outliers) pass (i.e greater than) the threshold $\hat{\Pi}_{thres}(r)$, so at most $r$ proportion of these pairs are false positives. If not (i.e. median-IQR/2 is not greater than $\hat{\Pi}_{thres}(r)$), $\alpha$ is too stringent with overall too low selection frequency, the previous $\alpha$ value will be the optimal one we want to use. If yes, we will continue to select a smaller $\alpha$ and repeat the above steps.

---

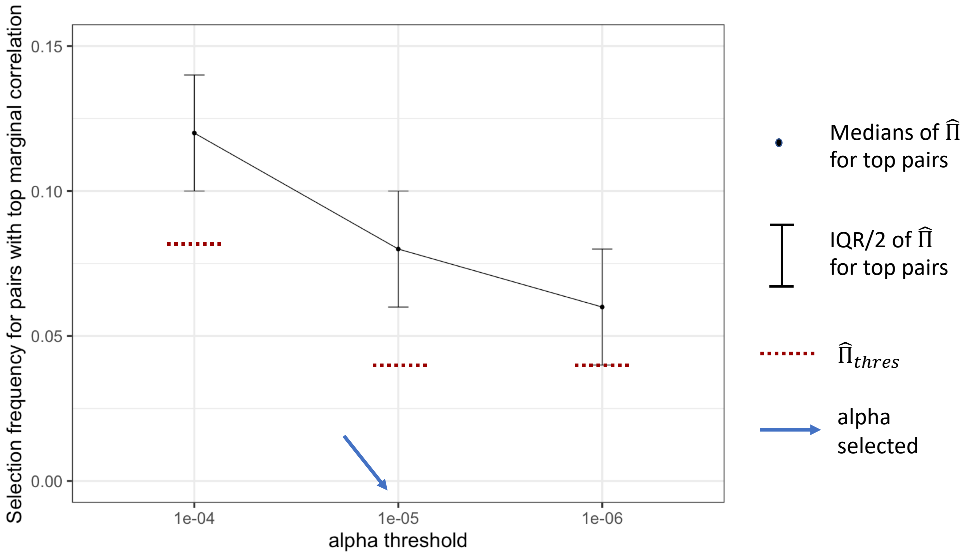


Figure S1: For pairs with top marginal correlations, compare $\hat{\Pi}_{(\phi)}$ to $\hat{\Pi}_{thres}(r = 0.1)$ for different $\alpha$ threshold in GBM data example. When $\alpha$ decreases to $1e - 6$, we see that the selection frequencies of top pairs no longer pass (i.e. not greater than) the selection frequency threshold $\hat{\Pi}_{thres}(r)$. Thus, $\alpha = 1e - 5$ will be selected in this case.

### 6 More results for simulation studies

#### 6.1 Scenario IA, $(\rho_X, \rho_Y) = (0.2, 0.8)$

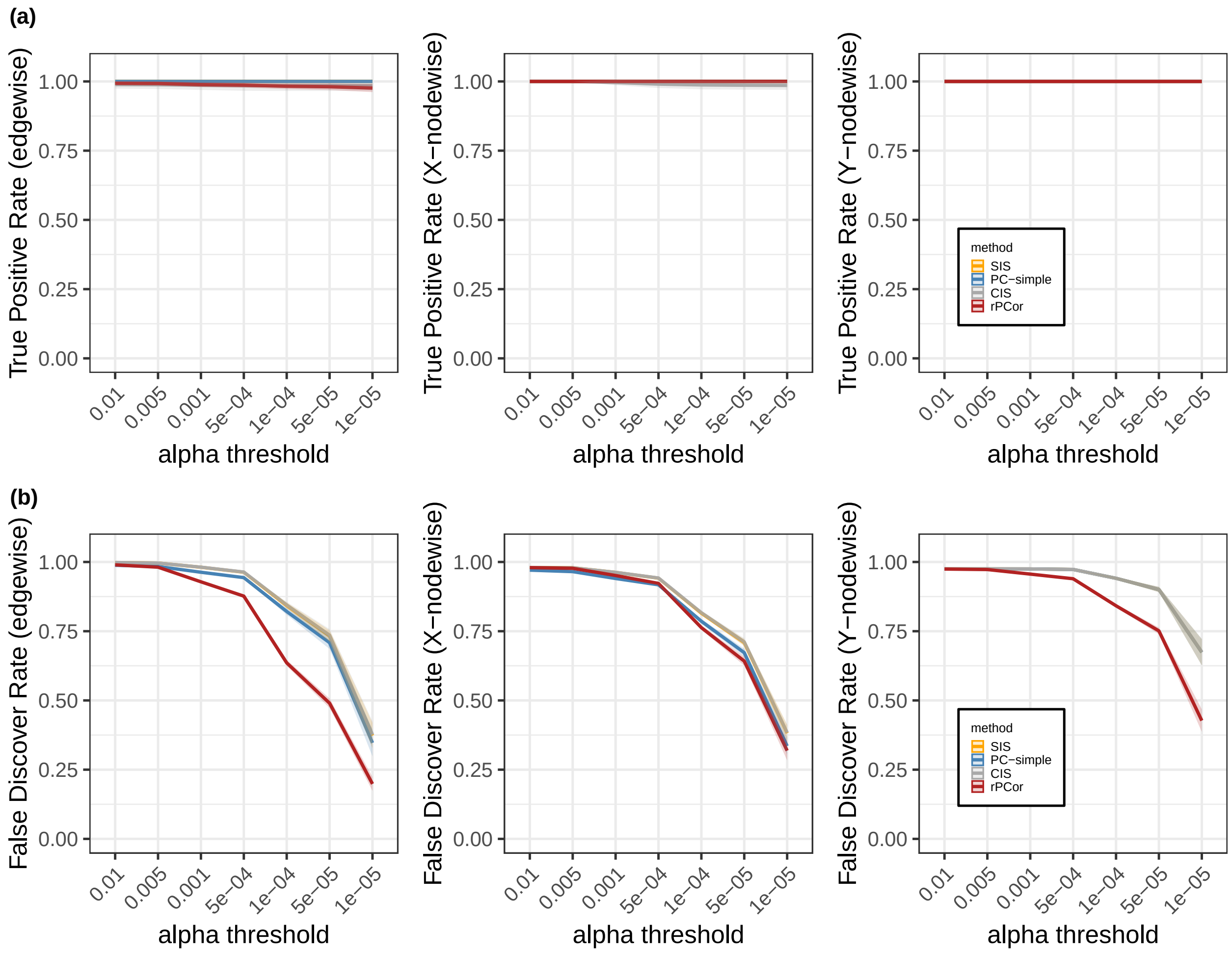


Figure S2: Simulation results for Scenario IA with $(\rho_X, \rho_Y) = (0.2, 0.8)$. (a) True positive rates comparison for edgewise, X-nodewise and Y-nodewise results (from left to right). (b) False discover rates comparison for edgewise, X-nodewise and Y-nodewise results (from left to right)

#### 6.2 Scenario IA, $(\rho_X, \rho_Y) = (0.8, 0.2)$

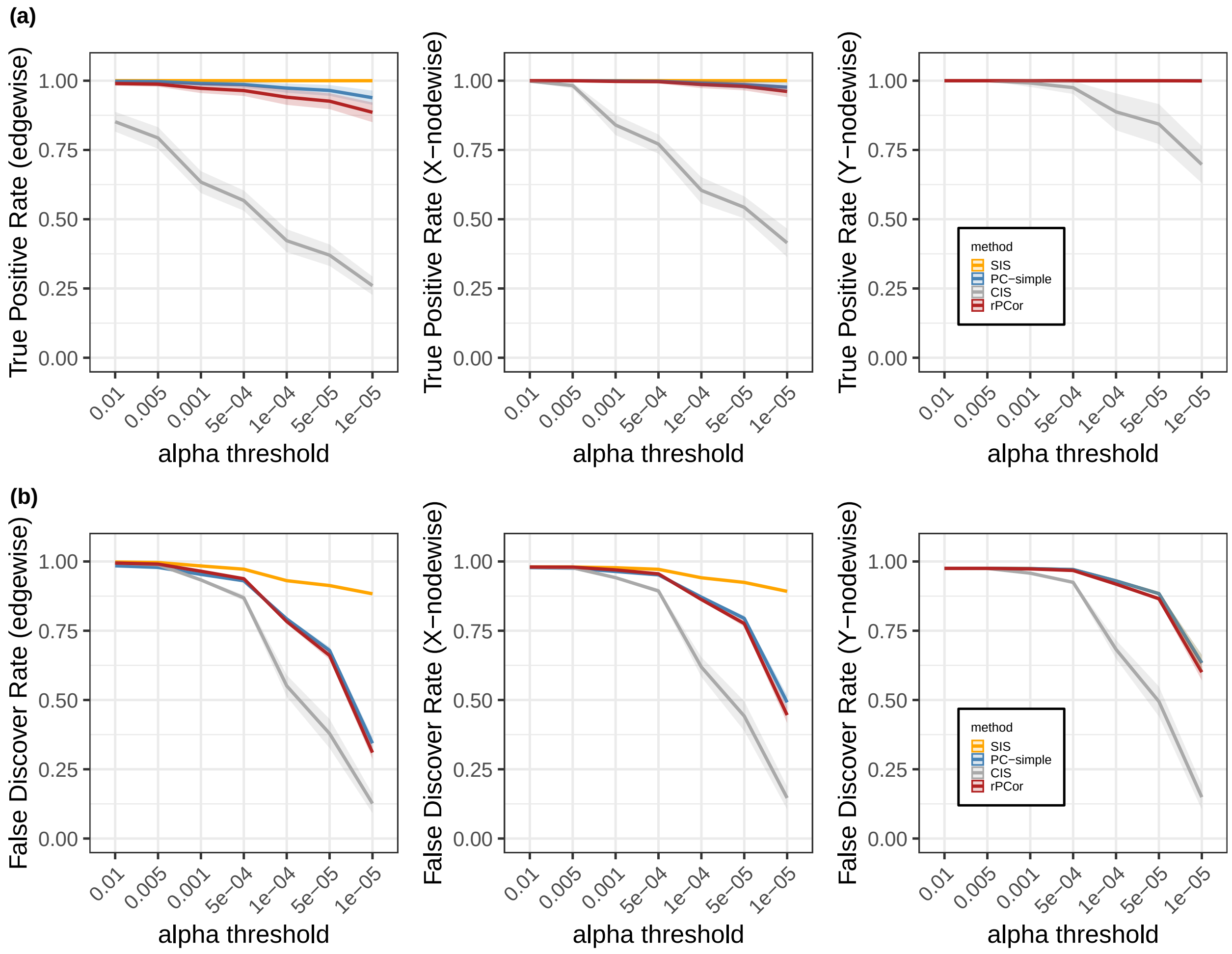


Figure S3: Simulation results for Scenario IA with $(\rho_X, \rho_Y) = (0.8, 0.2)$. (a) True positive rates comparison for edgewise, X-nodewise and Y-nodewise results (from left to right). (b) False discover rates comparison for edgewise, X-nodewise and Y-nodewise results (from left to right)

##### 6.3 Scenario IA, $(\rho_X, \rho_Y) = (0.2, 0.2)$

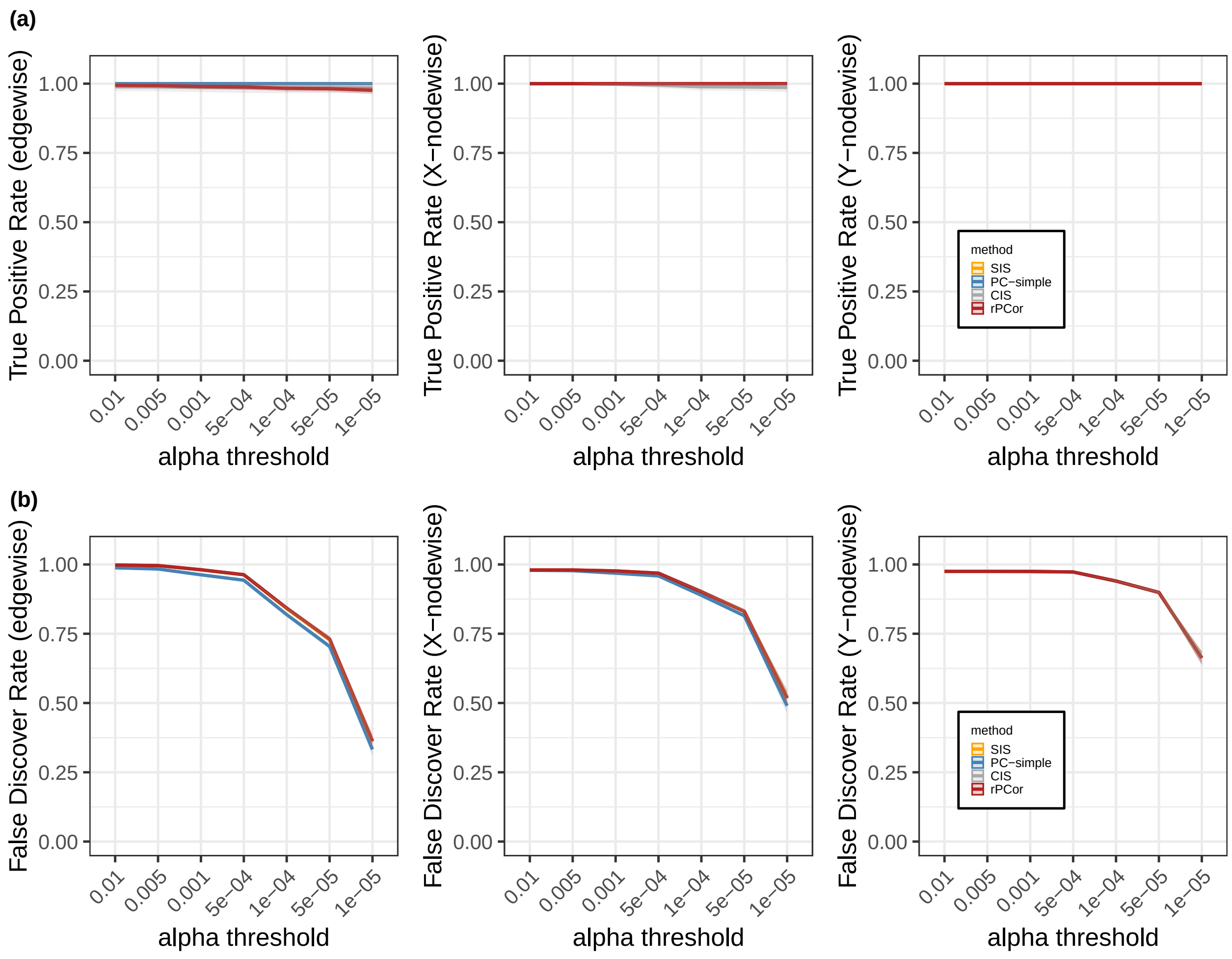


Figure S4: Simulation results for Scenario IA with $(\rho_X, \rho_Y) = (0.2, 0.2)$. (a) True positive rates comparison for edgewise, X-nodewise and Y-nodewise results (from left to right). (b) False discover rates comparison for edgewise, X-nodewise and Y-nodewise results (from left to right)

#### 6.4 Scenario IB

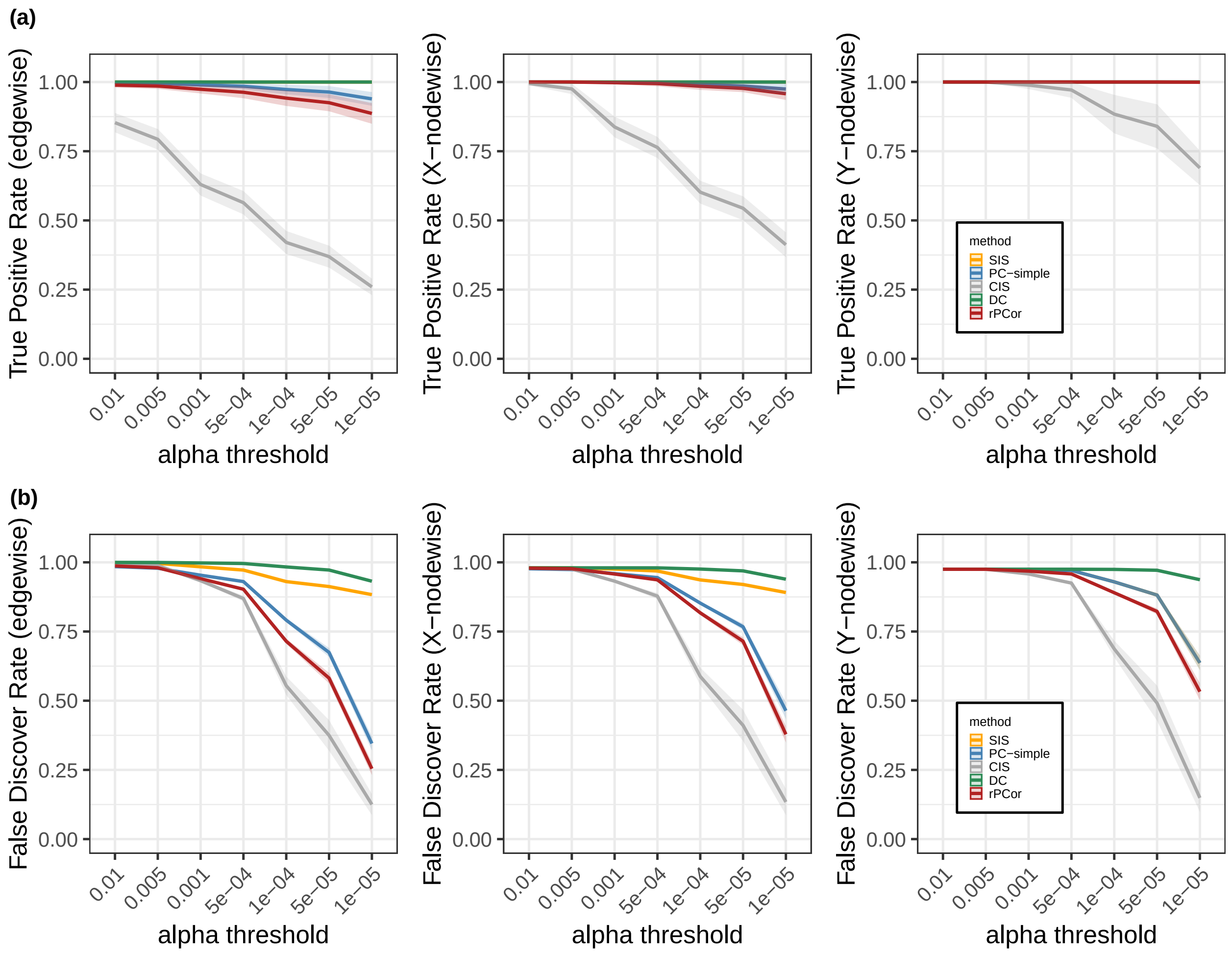


Figure S5: Simulation results for Scenario IB with $(\rho_X, \rho_Y) = (0.8, 0.8)$. (a) True positive rates comparison for edgewise, X-nodewise and Y-nodewise results (from left to right). (b) False discover rates comparison for edgewise, X-nodewise and Y-nodewise results (from left to right)

#### 6.5 Scenario II

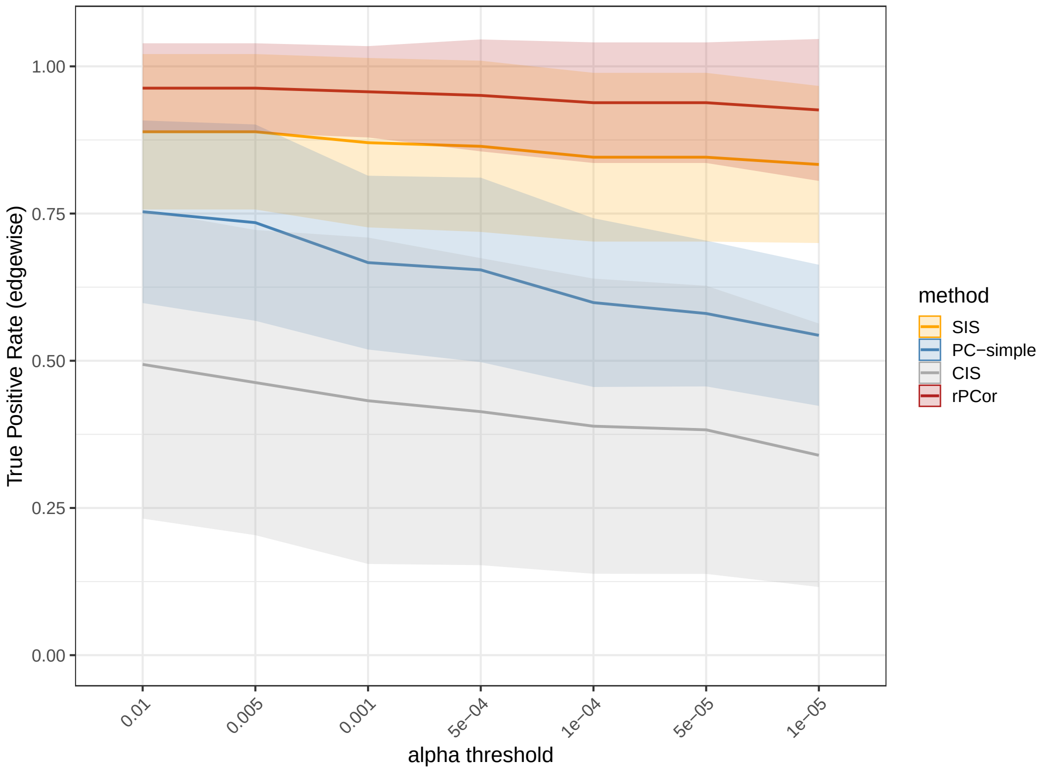


Figure S6: Simulation results for Scenario II comparing rPCor with the non-robust SIS, CIS, PC-simple methods in the presence of heavy-tailed data.

#### 6.6 Computation time comparison in simulation studies

Table S1: Computation time per replication for different scenarios and $(\rho_X, \rho_Y)$ (in seconds, mean and standard deviation calculated over 10 replication)

| | $(\rho_X, \rho_Y) = (0.8, 0.8)$ | |
|---|---|---|
| | Scenario IA | Scenario IB |
| SIS | 39.3 (8.9) | 40.7 (14.2) |
| PC-simple | 21231.4 (2521.3) | 21994.2 (3072.3) |
| CIS | 1239.4 (85.2) | 1259.5 (124.6) |
| DC-SIS | $>$ 3 days | $>$ 3 days |
| rPCor | 47.3 (3.5) | 51 (15.3) |
| rPcor-module | 46.1 (3.1) | 50.4 (15.1) |
| | $(\rho_X, \rho_Y) = (0.8, 0.2)$ | |
| | Scenario IA | Scenario IB |
| SIS | 49.3 (24.9) | 45.8 (21.9) |
| PC-simple | 22937.2 (4310.1) | 22281.7 (2470.2) |
| CIS | 1422.8 (93.7) | 1254.3 (275.9) |
| DC-SIS | $>$ 3 days | $>$ 3 days |
| rPCor | 60.7 (22.4) | 57.6 (18.5) |
| rPcor-module | 58.4 (25) | 56 (16.2) |

### 7 Downstream analysis results on the final identified features for Pan-kidney lncRNA regulation example

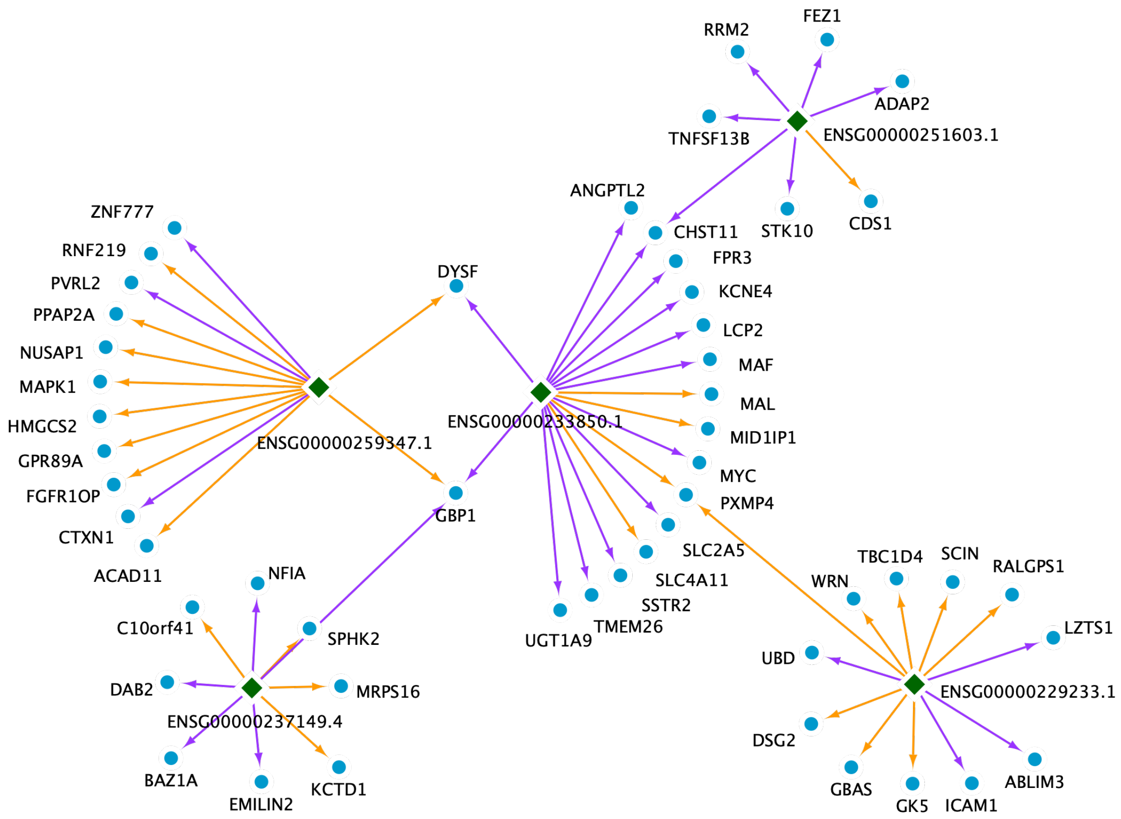


Figure S7: TCGA Pan-kidney lncRNA-gene regulation network example 1. Green colored nodes with diamond shape in the hub indicate the lncRNAs. Blue colored nodes with circular shape indicate the genes. Arrows in orange indicate positive regulation while arrows in purple indicate negative regulation (based on the estimated effect direction; see Table S7).

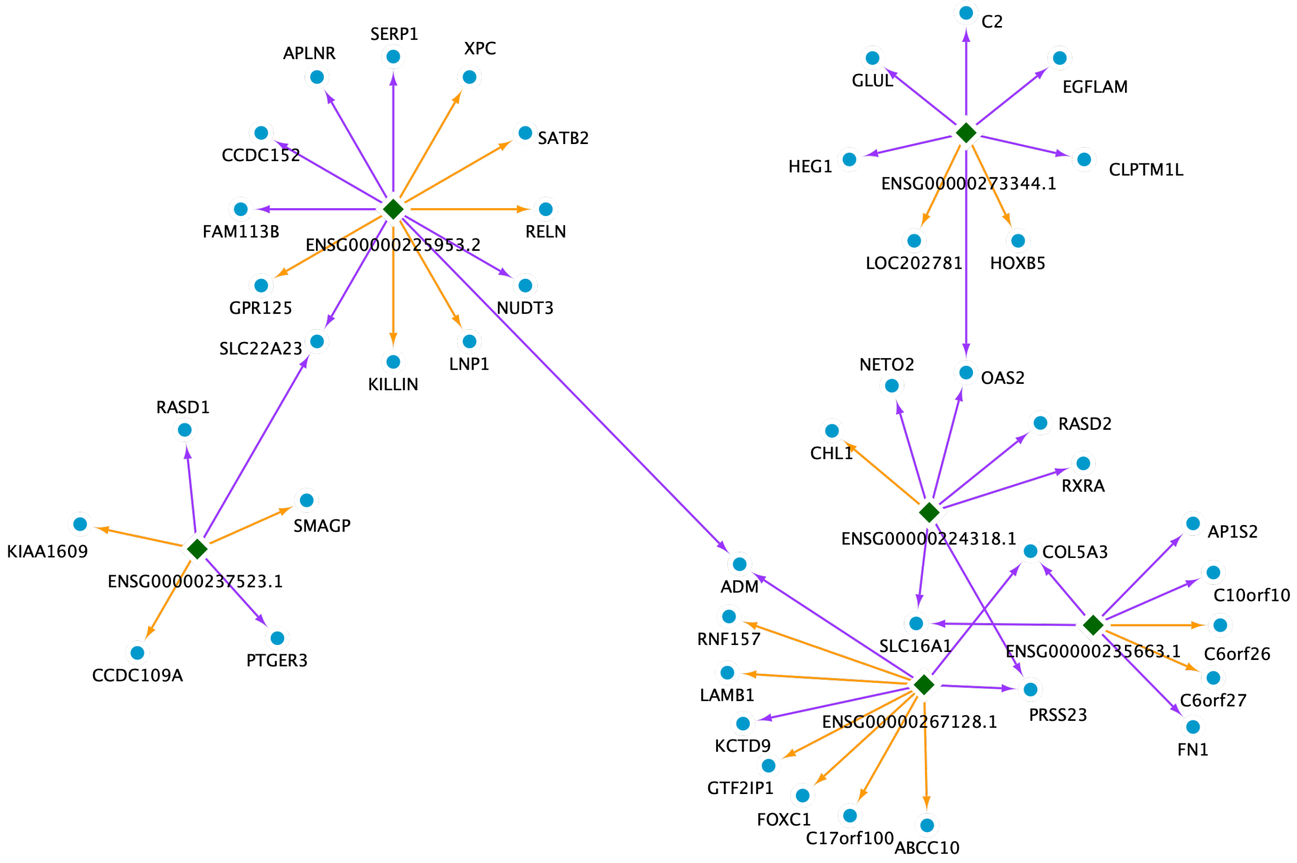


Figure S8: TCGA Pan-kidney lncRNA-gene regulation network example 2. Green colored nodes with diamond shape in the hub indicate the lncRNAs. Blue colored nodes with circular shape indicate the genes. Arrows in orange indicate positive regulation while arrows in purple indicate negative regulation (based on the estimated effect direction; see Table S7).

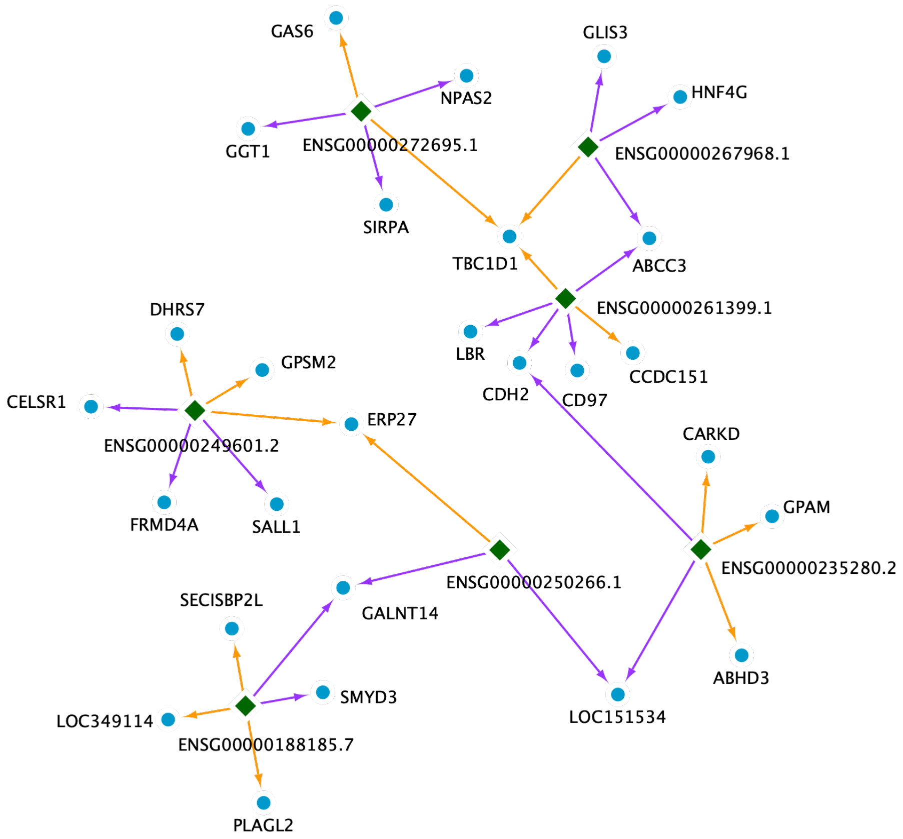


Figure S9: TCGA Pan-kidney lncRNA-gene regulation network example 3. Green colored nodes with diamond shape in the hub indicate the lncRNAs. Blue colored nodes with circular shape indicate the genes. Arrows in orange indicate positive regulation while arrows in purple indicate negative regulation (based on the estimated effect direction; see Table S7).

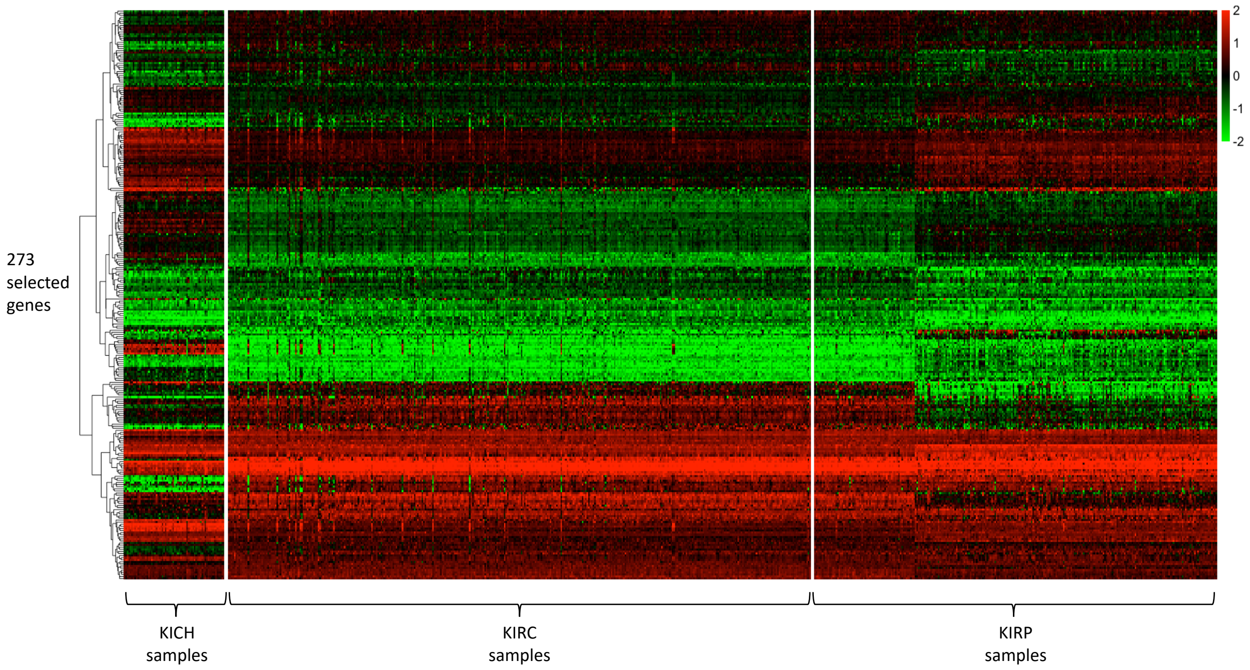


Figure S10: The heatmap of gene expression for 273 genes selected by our method in TCGA Pan-kidney example. Each row represents a gene and each column represents a sample. Samples from three different subtypes are separated. Genes are standardized and any values greater than 2 (or less than -2) are fixed as 2 (or -2).

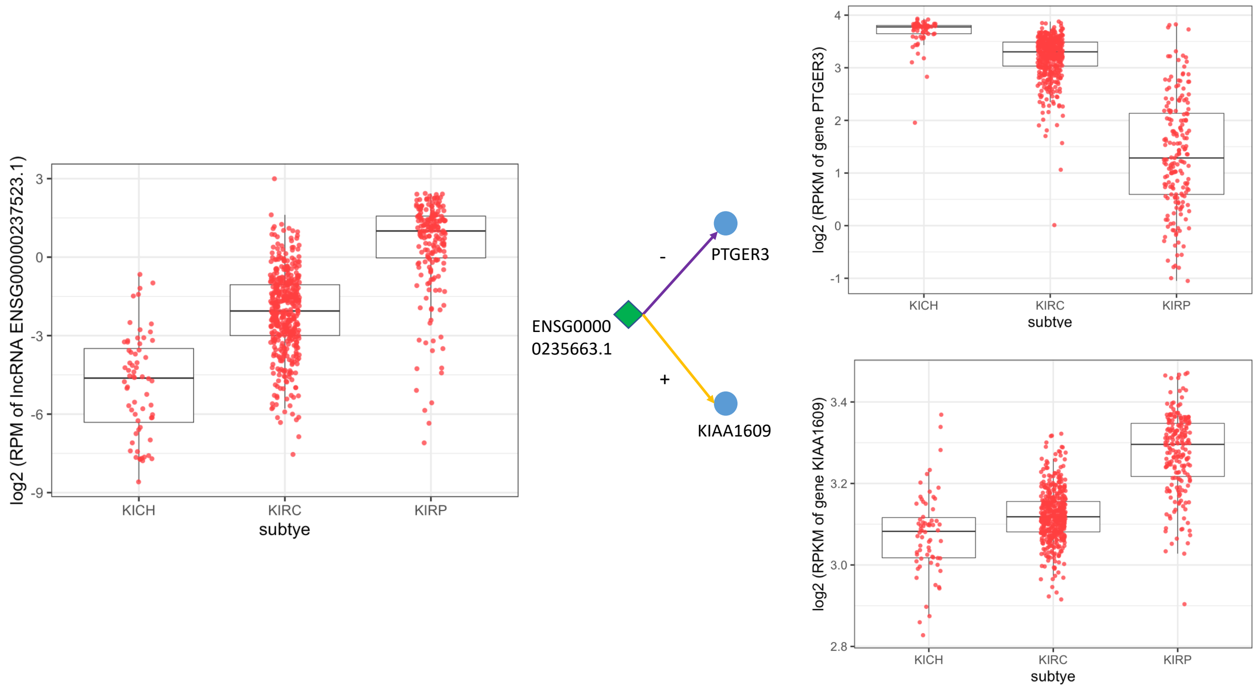


Figure S11: The association between cancer subtypes and the identified lncRNAs/genes in TCGA Pan-kidney example. Here we show a example of lncRNA (ENSG000000237523.1) and two of its target genes (PTGER3 and KIAA1609). Based on our estimation results, PTGER3 is being down-regulated while KIAA1609 is being up-regulated. The boxplots show that the different expression patterns of these lncRNA and genes in different Pan-kidney cancer subtypes.

### 8 Data preprocessing and results for GBM DNA methylation regulation example

#### 8.1 Data preprocessing

We downloaded the DNA methylation data measured by HM27K array (including about $27,000$ CpG sites, in zero-centered beta values) and the gene expression data measured by RNA-seq (in RPKM) of TCGA-GBM cohort from LinkedOmics (Vasaikar et al., 2018). The analysis was restricted to $n = 278$ GBM patients who have both DNA methylation and gene expression data available. We carefully preprocessed the data and followed a common practice by removing methylation sites with any missing values and those with less than 5% or more than 95% of samples having beta values less than 0.5 (Wang et al., 2013; Brennan et al., 2013). For the gene expression data, we only kept genes with mean RPKM values greater than 5. The processed data included $p = 6427$ CpG sites and $q = 8196$ genes for $n = 278$ GBM patients.

#### 8.2 Comparison results

Table S2: Screening and regularization results of different methods for GBM example

| method | # of pairs left | # of CpG sites left | # of genes left |
|---|---|---|---|
| orginal data | 52675692 | 6427 | 8196 |
| SIS | 1517203 | 6121 | 7860 |
| DC-SIS | 3796961 | 6408 | 8175 |
| PC-simple | 10495 | 966 | 7860 |
| rPCor | 3180 | 1870 | 2064 |
| rPCor + regularization | 481 | 180 | 429 |

#### 8.3 Downstream analysis results on the final identified features

| | | Transcription Factor: Yes | Transcription Factor: No | Total |
|---|---|---|---|---|
| Selected by our method | Yes | 42 | 387 | 429 |
| | No | 516 | 7251 | 7767 |
| | Total | 558 | 7638 | 8196 |

Table S3: Contingency table that shows the enrichment of Transcription Factors (TFs) among the genes selected by our method. The number of total genes (8196) is up to filtering. The p-value for Fisher's exact test is 0.0175.

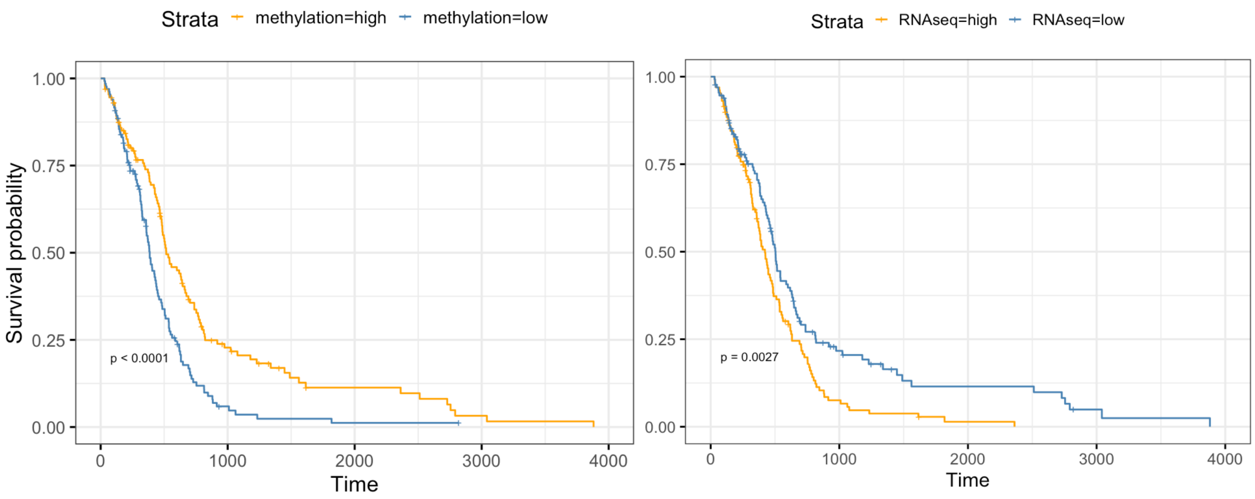


For the methylation site "cg07952391"    For the predicted target gene “THNSL2”

Figure S12: The association between survival time and the predicted methylation site/gene in TCGA GBM real data. Here the CpG site ”cg07952391” and gene ”THNSL2” is a CpG-gene regulatory pair identified by our method and their relationship is down-regulated. The Kaplan Meier Curves show that both of them are associated with survival time, but in an opposite direction.

Table S4: The list of TFs and their methylated CpG site regulators (in brackets) identified in our final model. Cis-regulated TF means that the CpG site is located in the promoter region of TF, and trans-regulated TF means that the CpG site is far away from the promoter region of TF, as defined by TCGA.

| | |
|---|---|
| cis-regulated TFs | AEBP1(cg02126753), HOXB2(cg09313705), NR2F2(cg08370996), SOX10(cg19257200), SP100(cg23539753) |
| trans-regulated TFs | ASCL1(cg14912034), BCL11A(cg01015871,cg02751839), BNC2(cg06454084), CC2D1A(cg07084163), CEBPB(cg08251399), CENPT(cg11976166), CREM(cg20200335), CTCF(cg12558519), CXXC4(cg09837648), EN1(cg11081833), ESRRA(cg22721827), HOXB7(cg07823492), HOXC10(cg12683929), MEOX2(cg15582789), MITF(cg06630241), MNX1(cg23173910), NKX2-5(cg18972811), OLIG2(cg26134665), POU6F1(cg02751839), PPARD(cg10011623), SAFB(cg26898336), SALL2(cg00622552), SHOX2(cg21416237), SIX1(cg09432376), SLC2A4RG(cg05668853), SOX4(cg09432376), TGIF2(cg21416237), TRPS1(cg17124700), TWIST1(cg10011623), ZBTB39(cg16483916), ZBTB5(cg19257550), ZNF212(cg15741583), ZNF219(cg22708914), ZNF248(cg07977490,cg08090640,cg09637363), ZNF358(cg16483916), ZNF419(cg13802966), ZNF423(cg09432376) |

### 9 Other supplementary tables

Table S5: Univariate association analysis of the identified 46 lncRNAs and 273 genes with the subtype (ANOVA model), pathological stage (linear regression) as well as survival time (Cox model) in the Pan-kidney example. P-values and Benjamini and Hochberg procedure (Benjamini and Hochberg, 1995) adjusted q-values are reported.

Table S6: Univariate association analysis of the identified 180 methylated CpG sites and 429 genes with the survival time (Cox model) in the GBM example. P-values and Benjamini and Hochberg procedure (Benjamini and Hochberg, 1995) adjusted q-values are reported.

Table S7: Final regression coefficient estimate $\hat{\beta}_{final}$ of the identified 296 lncRNA-gene pairs and 45 CpG-TF pairs in the Pan-kidney and GBM examples.

#### References


Benjamini, Y. and Hochberg, Y. (1995). Controlling the false discovery rate: a practical and powerful approach to multiple testing. *Journal of the Royal statistical society: series B (Methodological)*, 57(1):289–300.

Brennan, C. W., Verhaak, R. G., McKenna, A., Campos, B., Noushmehr, H., Salama, S. R., Zheng, S., Chakravarty, D., Sanborn, J. Z., Berman, S. H., et al. (2013). The somatic genomic landscape of glioblastoma. *Cell*, 155(2):462–477.

Bühlmann, P., Kalisch, M., and Maathuis, M. H. (2010). Variable selection in high-dimensional linear models: partially faithful distributions and the pc-simple algorithm. *Biometrika*, 97(2):261–278.

Chiquet, J., Mary-Huard, T., and Robin, S. (2017). Structured regularization for conditional gaussian graphical models. *Statistics and Computing*, 27(3):789–804.

Efron, B. (2012). *Large-scale inference: empirical Bayes methods for estimation, testing, and prediction*, volume 1. Cambridge University Press.

Even, S. (2011). *Graph algorithms.* Cambridge University Press.

Fan, J. and Lv, J. (2008). Sure independence screening for ultrahigh dimensional feature space. *Journal of the Royal Statistical Society: Series B (Statistical Methodology)*, 70(5):849–911.

He, K., Kang, J., Hong, H. G., Zhu, J., Li, Y., Lin, H., Xu, H., and Li, Y. (2019). Covariance-insured screening. *Computational statistics & data analysis*, 132:100–114.

He, K., Li, Y., Zhu, J., Liu, H., Lee, J. E., Amos, C. I., Hyslop, T., Jin, J., Lin, H., Wei, Q., et al. (2016). Component-wise gradient boosting and false discovery control in survival analysis with high-dimensional covariates. *Bioinformatics*, 32(1):50–57.

Ke, Y., Minsker, S., Ren, Z., Sun, Q., and Zhou, W.-X. (2019). User-friendly covariance estimation for heavy-tailed distributions. *Statistical Science*, 34(3):454–471.

Langfelder, P. and Horvath, S. (2008). Wgcna: an r package for weighted correlation network analysis. *BMC bioinformatics*, 9(1):1–13.

Li, R., Zhong, W., and Zhu, L. (2012). Feature screening via distance correlation learning. *Journal of the American Statistical Association*, 107(499):1129–1139.

Mazumder, R. and Hastie, T. (2012). Exact covariance thresholding into connected components for large-scale graphical lasso. *The Journal of Machine Learning Research*, 13(1):781–794.

Meinshausen, N. and Bühlmann, P. (2010). Stability selection. *Journal of the Royal Statistical Society: Series B (Statistical Methodology)*, 72(4):417–473.

Sohn, K.-A. and Kim, S. (2012). Joint estimation of structured sparsity and output structure in multiple-output regression via inverse-covariance regularization. In *Artificial Intelligence and Statistics*, pages 1081–1089. PMLR.

Székely, G. J., Rizzo, M. L., and Bakirov, N. K. (2007). Measuring and testing dependence by correlation of distances. *The annals of statistics*, 35(6):2769–2794.

Van Dam, S., Vosa, U., van der Graaf, A., Franke, L., and de Magalhaes, J. P. (2018). Gene co-expression analysis for functional classification and gene–disease predictions. *Briefings in bioinformatics*, 19(4):575–592.

Vasaikar, S. V., Straub, P., Wang, J., and Zhang, B. (2018). Linkedomics: analyzing multi-omics data within and across 32 cancer types. *Nucleic acids research*, 46(D1):D956–D963.

Wang, L., Zheng, C., Zhou, W., and Zhou, W.-X. (2020). A new principle for tuning-free huber regression. *Statistica Sinica*.

Wang, W., Baladandayuthapani, V., Morris, J. S., Broom, B. M., Manyam, G., and Do, K.-A. (2013). ibag: integrative bayesian analysis of high-dimensional multiplatform genomics data. *Bioinformatics*, 29(2):149–159.

Witten, D. M., Friedman, J. H., and Simon, N. (2011). New insights and faster computations for the graphical lasso. *Journal of Computational and Graphical Statistics*, 20(4):892–900.

Witten, D. M. and Tibshirani, R. (2009). Covariance-regularized regression and classification for high dimensional problems. *Journal of the Royal Statistical Society: Series B (Statistical Methodology)*, 71(3):615–636.
